# A model for *cis*-regulation of transcriptional condensates and gene expression by proximal lncRNAs

**DOI:** 10.1101/2023.01.04.522753

**Authors:** Pradeep Natarajan, Krishna Shrinivas, Arup K. Chakraborty

## Abstract

Long non-coding RNAs (lncRNAs) perform several important functions in cells including cis-regulation of transcription. Barring a few specific cases, the mechanisms underlying transcriptional regulation by lncRNAs remain poorly understood. Transcriptional proteins can form condensates via phase separation at protein-binding loci (BL) on the genome (e.g., enhancers and promoters). lncRNA-coding genes are present at loci in close genomic proximity of these BL and these RNAs can interact with transcriptional proteins via attractive heterotypic interactions mediated by their net charge. Motivated by these observations, we propose that lncRNAs can dynamically regulate transcription in cis via charge-based heterotypic interactions with transcriptional proteins in condensates. To study the consequences of this mechanism, we developed and studied a dynamical phase-field model. We find that proximal lncRNAs can promote condensate formation at the BL. Vicinally localized lncRNA can migrate to the BL to attract more protein because of favorable interaction free energies. However, increasing the distance beyond a threshold leads to a sharp decrease in protein recruitment to the BL. This finding could potentially explain why genomic distances between lncRNA-coding genes and protein-coding genes are conserved across metazoans. Finally, our model predicts that lncRNA transcription can fine-tune transcription from neighboring condensate-controlled genes, repressing transcription from highly expressed genes and enhancing transcription of genes expressed at a low level. This non-equilibrium effect can reconcile conflicting reports that lncRNAs can enhance or repress transcription from proximal genes.

**SIGNIFICANCE:** Long non-coding RNAs (lncRNAs) form a significant part of the human genome but do not code for any proteins. They have many hypothesized functions in the cell, including the regulation of transcription. Transcriptional condensates are assemblies of transcriptional proteins that concentrate at specific genomic sites through phase separation and can regulate transcription. In this study, we propose that lncRNAs can regulate transcription by interacting with proteins in transcriptional condensates to modulate condensate formation. We find that this model can explain some puzzling observations such as conflicting reports of gene activation and repression by lncRNAs, and conservation of genomic distances between lncRNA-coding genes relative to protein-coding genes in metazoans. Experimentally testable predictions that can further explore our model are discussed.

## INTRODUCTION

Genes that encode long non-coding RNAs (lncRNAs) outnumber protein-coding genes (PCGs) in the mammalian genome (1, 2). lncRNAs are RNAs that have a length of>200 nucleotides and are not translated into any proteins unlike the messenger RNAs (mRNAs). Some well-studied lncRNAs include NEAT1 which acts as a scaffold in paraspeckles, MALAT1 which regulates the phosphorylation of SR proteins in nuclear speckles, XIST which is involved in the silencing of the X chromosome, and NORAD which promotes genomic stability (3). Except for these and a small number of others, the biological function of the vast majority of lncRNAs is poorly understood.

There is an emerging body of literature that suggests that lncRNAs can regulate transcription in *cis* (4–9). lncRNAs involved in *cis*-regulation usually affect transcription in a manner that depends on their genomic locus. Transcription of these lncRNAs has a local effect and directly correlates with the transcription of PCGs in genomic and spatial proximity in most cases (10–12). However, recent experiments that perturb lncRNA transcription report conflicting observations on its impact on transcription from neighboring genes. Luo and coworkers knocked down several divergent lncRNAs in mouse embryonic stem cells using RNAi and observed that gene expression from neighboring PCGs went up in some cases while it went down in others (4). Engreitz and coworkers suppressed lncRNA transcription in mouse cell lines by knocking out their promoters and reported a similar observation (5). The promoter knockout in some rare cases dramatically decreased gene expression from the neighboring PCG. We do not have a unifying framework to explain these seemingly conflicting observations.

Several experimental studies offer a glimpse into the mechanisms by which lncRNAs regulate transcription in *cis*. lncRNAs can activate gene expression by recruiting the transcriptional coactivator Mediator to neighbor genes (4, 9), promote looping between enhancers and promoters (4, 8) and recruit histone modifiers to promoter regions of neighboring genes (7). The process of lncRNA transcription has also been hypothesized to activate the transcription of target genes by maintaining enhancers in an active state (13) and by increasing the local concentration of transcription-associated proteins at neighboring promoters (5). The cis-regulatory function of lncRNA sequences does not appear to depend strongly on their specific sequences as they are often poorly conserved (6, 14) and only weakly selected in humans (15). However, recent evidence suggests that lncRNAs occur at conserved genomic positions relative to orthologous genes (6, 16, 17). This kind of “positional” conservation rather than sequence conservation motivated us to consider a physical mechanism for *cis*-regulation of gene expression that is agnostic to the specific lncRNA sequence.

Using RNA-DNA SPRITE, Quinodoz and coworkers demonstrated that mature lncRNAs tend to localize in the vicinity of their coding genomic regions and form their own compartments (18). There is emerging evidence that transcriptional proteins also form their own compartments – called transcriptional condensates – at enhancers and promoters (19–23) and control gene expression from target genes (24, 25). These condensates are comprised of biomolecules including transcription factors (19), transcriptional coactivators (20, 23), and RNA Polymerase II (22, 23) that are recruited to enhancers and promoters via a phase-separation mechanism (26). Promoters of PCGs are surrounded mostly by lncRNA-coding genes in their immediate genomic and spatial neighborhood (4, 10, 11) and many enhancer loci also code for lncRNAs (27, 28). The spatial distance between lncRNA-coding loci and promoters and enhancers is of the same order as the size of stable transcriptional condensates (Refer to section S1 in the supplemental material). Given this spatial proximity, lncRNAs could interact with components of the transcriptional condensate.

Motivated by these observations, we hypothesized that lncRNAs can regulate transcription in *cis* by interacting with the components of the transcriptional condensate. But what is the nature of this interaction? Recent work suggests that transcriptional coactivators such as Mediator subunit 1 and BRD4 have positively charged disordered domains that can interact with the negatively charged RNA polymer (29) via screened electrostatic interactions. This can result in the condensation of transcriptional proteins driven by the phenomenon of complex coacervation (30–32). A small concentration of RNA promotes condensation driven by electrostatic attraction between the differently charged polymers. However, when the RNA concentrations exceed a value that corresponds to a balance between the total positive and negative charge in the system, this leads to condensate dissolution driven by entropic effects of confining the polymer within the coacervate and electrostatic repulsion between like-charged RNAs (33, 34). The non-equilibrium process of RNA transcription can therefore feedback on itself by initially aiding condensate formation and then dissolving it (29). This provides a sequence-agnostic biophysical mechanism that could also be employed by many lncRNAs to control transcription in *cis*.

In this paper, we study how lncRNAs may regulate transcriptional condensates via non-equilibrium phenomena coupled to complex coacervation. We develop a phase-field model for transcriptional regulation by lncRNAs that incorporates known observations about lncRNAs, transcriptional condensates, and interactions between their components, and numerically simulate the model equations. Using this model, we predict that vicinally localized lncRNAs can reduce the threshold protein concentrations required for transcriptional condensate formation and increase protein recruitment to protein-binding loci (BL) on chromatin (e.g. enhancers and promoters). This is a local effect and drops off sharply with the distance between the lncRNA locus and the BL. Finally, we also predict that local transcription of lncRNAs can aid the formation of transcriptional condensates at PCGs or dissolve it, depending on their level of expression. This in turn has a corresponding effect on transcription from the PCGs. We predict that transcription of proximal lncRNAs enhances transcription from PCGs expressed at a low level, while the same process represses transcription from highly expressed PCGs. Based on these results, we propose that lncRNA transcription can act as a regulatory knob to fine-tune transcription from neighboring genes. Our model provides a mechanistic framework that reconciles conflicting observations about *cis*-regulation of transcription by lncRNAs, provides a possible explanation for how this function can impose genomic constraints on the positions of lncRNA loci and makes predictions that can be experimentally tested to further explore this mechanism.

## MODEL DESCRIPTION

We adopt a continuum phase-field approach to build our model. We have three biomolecular species in our model: the lncRNA, mRNA, and transcriptional proteins (Fig. 1A). We treat the latter as a quasi-species that includes all proteins related to the transcriptional machinery such as transcriptional coactivators and transcription factors. Each of these species is characterized by a concentration field - *ϕ_R_* (for lncRNA), *ϕ_M_* (for mRNA), and *ϕ_P_* (for protein) – which depends on the spatial position. These concentration fields evolve in time governed by partial differential equations (PDE) that describe (i) the transport of these species in space as a consequence of their interaction with each other and (ii) any reactions they might undergo.

**Figure 1:**
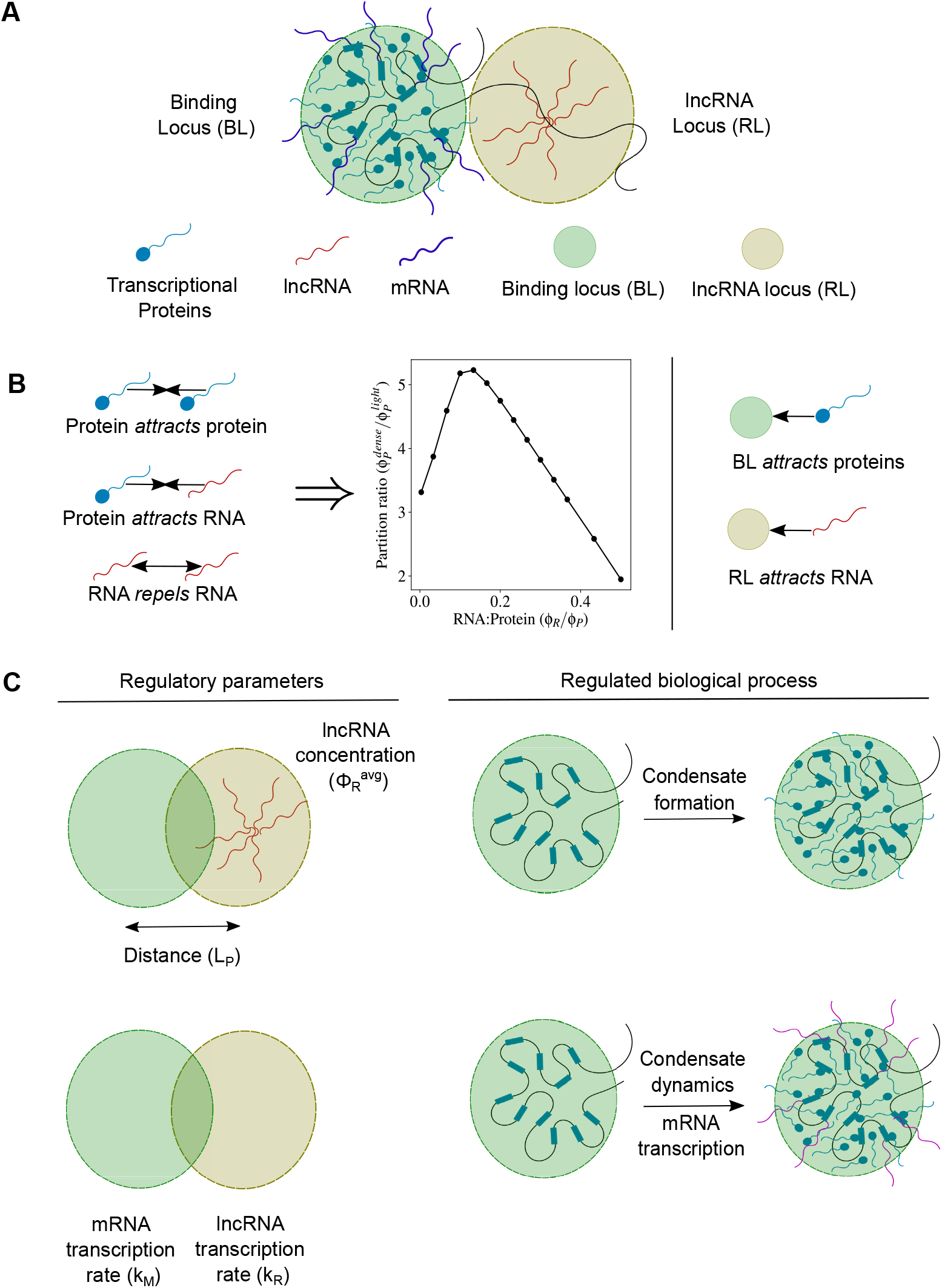
**A** Cartoon describing the molecular players involved in transcriptional condensate formation. **B** Transcriptional proteins attract each other through interactions mediated by their intrinsically disordered domains. RNAs (both lncRNA and mRNA) attract transcriptional proteins through interactions mediated by screened electrostatics or otherwise. RNAs repel each other due to electrostatic repulsion between like-charged polymers. These interactions result in re-entrant condensation of proteins. Transcriptional proteins can concentrate at chromatin regions rich in enhancers and promoters to form a dense phase which we call a transcriptional condensate. We call these regions of chromatin the binding locus (BL). lncRNAs localize near their genomic loci which we call the lncRNA locus (RL). **C** Cartoon describing the different regulatory parameters investigated in this study along with the biological process that they regulate. The amount of lncRNA (as measured by the average lncRNA concentration 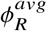) and the distance (*L_P_*) between the RL and the BL can affect condensate formation at the BL. The relative magnitudes of the mRNA transcription rate constant (*k_M_*) at the BL and the lncRNA transcription rate (*k_L_*) at the lncRNA locus affects the dynamics of the protein condensate and therefore mRNA transcription from the BL.

To account for the interactions between lncRNA, mRNA, protein, and the chromatin (summarized in Fig. 1B), we write down an expression for the free energy of this multi-component system that comprises the following three terms:

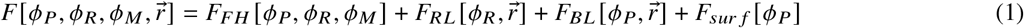

*F_FH_* [*ϕ_P_, ϕ_R_, ϕ_M_*] is a Flory-Huggins free energy that captures the self and cross interactions between transcriptional proteins, lncRNA, and the mRNA. A detailed expression for this free energy is given in section S2.3 of the supplemental material. The rationale behind choosing values for the different parameters associated with this expression is discussed in sections S2.1-S2.3 in the supplemental material, and the specific values used in simulations are summarized in supplementary table 1.

Briefly, this free energy captures the following three biologically relevant interactions: (i) attractive protein-protein interactions (ii) repulsive RNA-RNA interactions, and (iii) attractive protein-RNA interactions. We assign the protein-protein interactions to be attractive motivated by the observation that many transcriptional proteins contain intrinsically disordered regions (IDRs) that promote the formation of transcriptional condensates (19, 21). The attractive interactions between IDRs arise from various interactions at the amino acid level such as electrostatic (35, 36), pi-pi (37), cation-pi (38) and hydrophobic (35) interactions. Interactions between all the RNA species in our model are chosen to be repulsive motivated by the fact that lncRNA and mRNA species are both negatively charged polymers that can interact via screened electrostatic repulsion. Finally, the protein-mRNA and protein-lncRNA interactions are attractive in our model, motivated by the observation that many transcriptional coactivators contain positively charged IDRs (29) and transcription factors contain positively-charged RNA-binding regions (39) that can bind to negatively-charged RNA.

There is also emerging evidence that many lncRNAs localize in close proximity to their genomic loci (18). We refer to the genomic loci that code for lncRNAs as the lncRNA locus, or RL, for the rest of this paper. There are many mechanisms that could facilitate attractive interactions between lncRNAs and their RL – these include tethering by transcription factors such as YY1 (40, 41) or by RNA polymerase (42). Irrespective of the mechanism, we can write down a free energy between the lncRNA concentration field 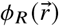 at position 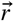 and its RL located at position 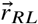 using a Gaussian function that has a range *σ_RL_* and strength of attraction *c_R_*:

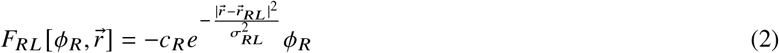

Finally, the term 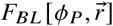 captures the interaction free energy of transcriptional proteins with regions of attractive chromatin that promote condensate formation (21). We call these regions of attractive chromatin such as specific enhancers, super-enhancers, or promoters as the binding locus, or BL. We can write down a free energy between transcriptional protein concentration field 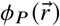 at position 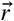 and its BL located at position 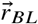 using a Gaussian function that has a range *σ_BL_* and strength of attraction *c_P_*:

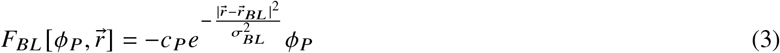

The term 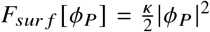 is a surface tension term that penalizes sharp gradients in protein concentration, with *κ* being the strength of this energy penalty. This term is not particularly important for our results but ensures that any phase separation is accompanied by smooth boundaries between phases.

Using this model, we hope to answer the following two questions: **(1) How does a lncRNA localized near a BL affect the formation of transcriptional condensates? (2) How does an actively transcribed lncRNA affect mRNA transcription from a nearby BL?** Specifically, we look at how the amount of lncRNA 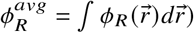, the distance between the BL and the RL 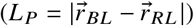, and the rate of lncRNA transcription at the RL (*k_R_*) relative to the mRNA (*k_M_*) affect the above processes.

### Dynamics of condensate formation

In this section, we develop a model to answer the first question: How does a lncRNA localized near a BL affect the formation of transcriptional condensates? To do this, we consider a situation where there is a uniform concentration of transcriptional proteins everywhere in space at time *t* = 0. Some amount of lncRNA is spatially localized at the RL which is present in the vicinity of the BL. There is no active transcription of mRNA happening at the BL and the free energy, in this case, does not depend on *ϕ_M_* i.e. 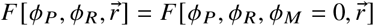. As time progresses, the protein starts to accumulate at the BL driven by the attractive protein-protein and protein BL interactions. We define a condensate as a region in space where the protein concentration is above a threshold value, which is set by the free energy parameters (Refer to section S2.4 in the supplemental material for details on this threshold value). The lncRNA localized at the RL can perturb the dynamics of condensate formation at the BL depending on its amount 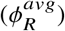 and how far away it is (*L_P_*).

Condensate formation happens over the time scale of a few minutes (24) which is much shorter compared to the half-lives of most lncRNAs (43) and proteins (44) which can span hours. Therefore, we assume that the protein and lncRNA are stable over our simulation of condensate formation. For conserved species, the spatiotemporal dynamics of concentrations are such that the molecules move down gradients in chemical potential. The coupled dynamics of the concentrations 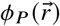 and 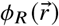 can be captured using the following Model B equations (45):

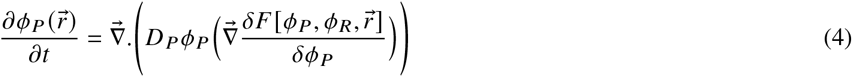

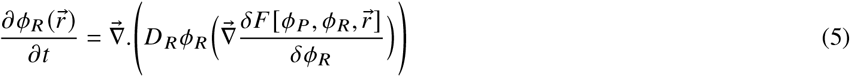

### Dynamics of transcription

In this section, we develop a model to answer the second question: How does an actively transcribed lncRNA affect mRNA transcription from a nearby BL? BLs with active mRNA transcription are often not isolated but located in neighborhoods that contain other actively transcribing RNAs including lncRNAs. Transcription of neighboring lncRNAs can potentially couple to the dynamics of mRNA transcription specifically by modulating protein recruitment to the BL and transcriptional condensate formation, thereby regulating gene expression.

Active transcription and depletion of RNAs that consume ATP can alter the local RNA concentrations and push the system far out of equilibrium. The rate of mRNA transcription must depend on both the local concentration of transcriptional proteins and the coding DNA. We take into account the local coding-DNA concentration through an effective rate constant that is a Gaussian function in space centered at the BL, reflecting the concentration of these genes at the BL. In addition to the spatially varying rate constant, the mRNA transcription rate has a simple first-order dependence on *ϕ_P_*, reflecting the activating effect of transcriptional proteins. To be general, we assume that lncRNA transcription is not controlled by the same transcriptional proteins and its rate is independent of *ϕ_P_*. The lncRNA transcription rate is also modeled as a Gaussian function in space centered at the RL, to reflect its transcription from its coding DNA which is localized at RL. Using this function for both the coding-DNA concentrations is a simple approximation if we assume the genomic region to be a Gaussian polymer. The values *σ_R_* and *σ_M_* reflect the spatial extents of the DNA that codes for the lncRNA and the mRNA respectively. In addition to the spatially varying production rates of the species, we also have a simple first-order decay of the lncRNA and mRNA species throughout space with rate constants of *k_dR_* and *k_dM_* respectively.

To understand how lncRNA transcription perturbs mRNA transcription, we use the following model where the reaction-diffusion dynamics of the lncRNA affect mRNA transcription by perturbing the dynamics of the protein field 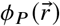:

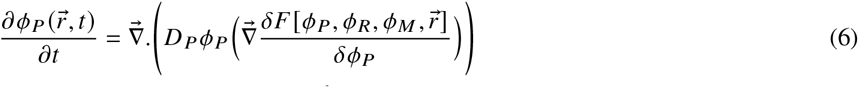

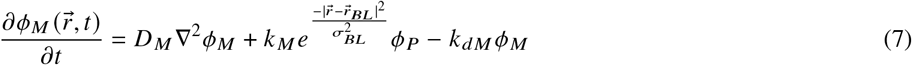

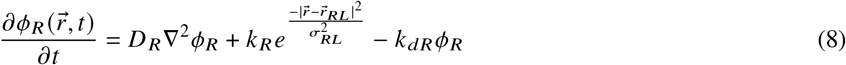

Section S2.6 of the supplemental material summarizes the specific values and ranges of parameters that were used for the simulation and a rationalization for these choices. For this study, we vary the magnitudes of the lncRNA production rate (*k_R_*) and mRNA transcription rate constant (*k_M_*), and investigate how that affects condensate dynamics and mRNA expression.

A difference between the model described by Eqs. 6-8 and that in the previous section is the mechanism of lncRNA localization. In this model, the lncRNA production rate is peaked at the RL. Therefore, the lncRNA concentration is highest at the RL and decreases with distance due to diffusion and degradation. Another important difference is that Eqs. 6–8 define processes far out of equilibrium, and not dynamics down a free energy gradient.

### Numerical simulation of model equations

The above partial differential equations were numerically solved using a custom python code, available here. The Zenodo-generated DOI for the same is 10.5281/zenodo.7461653. This code uses the finite volume solver Fipy developed by the National Institute of Standards and Technology (46). All simulations in this paper were done in a 2D circular domain of radius 15 units, with a circular discrete mesh. The spatially discretized PDEs were solved for each incremental time step with adaptive time stepping to pick smaller or larger time steps depending on how quickly or slowly the concentration fields change. A grid size of Δ*r* = 0.1 and a typical time step size on the scale of Δ*t* = 0.2 worked well for the simulations. Simulations were run for a duration of 2000 time steps, which was sufficient for the system to reach a steady state.

For the dynamics of condensate formation with localized lncRNA, the equilibrium concentration profile of lncRNA was obtained as described in section S2.2 of the supplemental material, which was then used as the initial condition for simulating the dynamics. For all simulations, a uniform protein concentration profile was used as the initial condition, with a value of 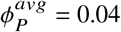 unless stated otherwise. This corresponds to a regime where the protein does not form a condensate by itself and requires lncRNA for condensate formation and this value was chosen to illustrate the effects of lncRNAs more sharply. The initial concentration of mRNA everywhere was set to *ϕ_M_* = 0. The no-flux Neumann boundary condition was applied to all species at the domain boundaries.

### Analyses

Numerical simulations yield the full concentration profiles of the protein 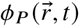, lncRNA 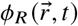, and mRNA 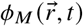 at all times *t*. Once we have this data, we can calculate quantities such as the concentration of a species at the BL, the partition ratio of species at the BL, the average concentration of the species in the system, and the chemical potential of the species. The precise formula for each of these quantities is described in section S2.8 of the supplementary material.

## RESULTS

### Proximal lncRNAs can enhance recruitment of transcriptional proteins to super-enhancers and promoters

Condensate formation by transcriptional proteins at BL is driven cooperatively by protein-chromatin binding interactions and attractive protein-protein interactions mediated by their disordered domains (21). When the concentration of transcriptional proteins crosses a threshold, there is a sharp increase in protein concentration at the BL due to phase separation and condensate formation driven by these two interactions.

As the first step, we wanted to understand how lncRNAs localized near a BL can affect condensate formation. The amount of lncRNA 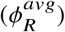 is an important regulatory parameter that controls the magnitude of this effect. We started with 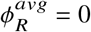 (no lncRNA) and progressively increased the amount of lncRNA in the system. We numerically simulated the model described by Eqs. 4-5 by varying the protein concentration in the system 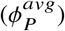 and quantified the protein partitioning to the BL at steady state (Fig. 2A). We find that vicinally localized lncRNAs consistently enhance protein partitioning to the BL compared to the base case where there is no lncRNA (Fig. 2B). Protein partitioning to the BL increases sharply upon increasing the protein concentration before reaching a plateau. This sharp increase is due to the phase separation of the proteins, and we can define a threshold value of protein concentration for which a condensate i.e. a dense phase of protein (with concentration ≥ 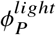) starts to appear at the BL. We find that lncRNAs localized near BL can reduce the transcriptional protein concentration thresholds that are required for phase separation and condensate formation (Fig. 2B). Thus, attractive interactions between transcriptional proteins and lncRNAs localized in the vicinity mediated by screened electrostatic interactions or otherwise can add an additional layer of cooperativity along with protein-chromatin and protein-protein interactions to aid condensate formation.

**Figure 2:**
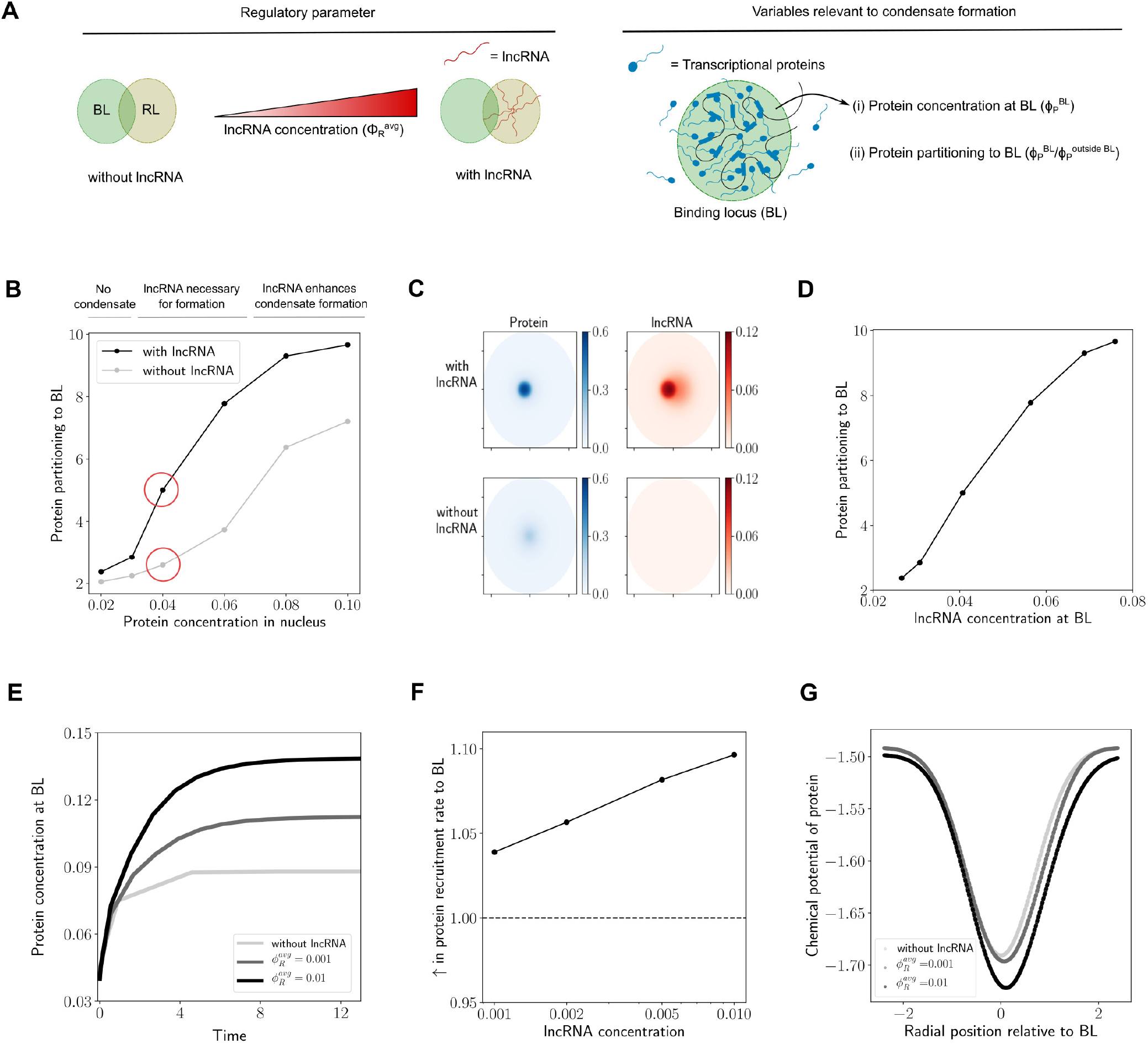
**A** In this figure, results are shown for what happens when we increase the lncRNA concentration 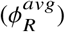 starting from a case without lncRNA 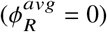. We quantify protein recruitment to the BL using the following two metrics: the protein concentration 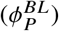 in the BL and the protein partitioning to the BL 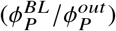. The distance between the loci was set to *L_P_*/*σ* = 0.8 **B** *Condensate formation:* Change in protein partitioning to the BL upon increasing the amount of protein in the nucleus. A protein condensate is formed when there is a sharp increase in protein partitioning to the BL. The grey curve corresponds to the case without lncRNA and the black curve corresponds to a case with a lncRNA amount of 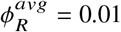. The concentration profiles of protein and lncRNA in space are depicted for the circled data points in figure C. **C** The protein and lncRNA concentration profiles are illustrated for the case with and without lncRNA. The average protein concentration in the nucleus for both cases is 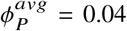. **D** The relationship between protein partitioning to the BL and the average lncRNA concentration in the BL for different amounts of protein in the nucleus. **E** *Dynamics of protein recruitment:* Protein concentration in the BL vs. time for different amounts of lncRNA. The time (*t*) is reported in dimensionless units as *tD_P_/R*^2^. *D_P_* is the diffusion coefficient of the protein and *R* is the radius of the nucleus. **F**The initial rate of protein recruitment to the BL for different amounts of lncRNA. The initial rate of protein recruitment is the slope of the graphs in figure E at *t* = 0. They are reported in this figure as a ratio relative to the case with no lncRNA 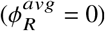. **G** Chemical potential of protein vs. radial position at *t* = 0 for different amounts of lncRNA. The radial position is measured relative to the center of the BL, with the origin being the center.

There exists a regime of protein concentrations for which lncRNA is necessary for condensate formation (Fig. 2B) and a condensate does not form in the absence of lncRNAs (Fig. 2C). In this regime, the additional layer of cooperativity added by the lncRNA-protein attractive interactions is necessary for condensate formation. This observation can explain why knocking down lncRNAs can sometimes have a dramatic effect on mRNA transcription from neighboring genes (5). A transcriptional condensate simply does not form to initiate transcription. At large protein concentrations where condensate formation happens even in the absence of lncRNAs, the presence of lncRNAs in the vicinity can still enhance protein recruitment to the BL (Fig. 2B). In all cases, protein partitioning to the BL directly correlates with the lncRNA concentration at the BL (Fig. 2D).

The dynamics of protein recruitment to the BL dictates the speed of cellular response to an external stimulus by activating gene expression. Therefore, we wanted to understand how different amounts of proximally localized lncRNA 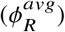 affect the dynamics of protein recruitment to the BL. We graphed the evolution of protein concentration at the BL with time (Fig. 2E) and find that increasing the amount of lncRNA has two distinct effects, which point to two distinct regulatory roles: (i) Higher amounts of lncRNA can increase the initial rate of protein recruitment to the BL (Fig. 2F), speeding up the response time between the cells receiving a stimulus and forming transcriptional condensates, (ii) Higher amounts of lncRNA can increase the protein concentration at the BL at steady state (Fig. 2E), increasing the strength of response to the stimulus. In this way, a cell can regulate the speed and magnitude of protein recruitment to the BL by using the amounts of proximally localized lncRNAs as a tunable knob.

To shed light on the mechanistic basis of these effects, we graphed the chemical potential profiles of the protein at initial times (Fig. 2G). The chemical potential at initial times has a shape of a Gaussian well, which is what we would expect based on the attractive protein-chromatin interactions at the BL described by Eq. 3. Increasing the amount of lncRNA 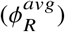 in the vicinity of the BL has two effects: it makes the well deeper and broader. The presence of lncRNAs near the BL and their attractive interactions with the protein provides a free energy benefit in addition to the protein-chromatin interactions, which translates to a deeper chemical potential well. A deeper well means that the chemical potential gradients are steeper, resulting in higher fluxes of the protein and a faster speed of protein recruitment to the BL. Spatial overlap between the BL and the localized lncRNA results in a broader effective region in space that attracts the protein. A broader well leads to increased overall protein recruitment to the BL as a broader well can hold more overall amount of protein.

In summary, the two ingredients – (i) localization of lncRNA near BL and (ii) attractive interactions between lncRNAs and proteins, possibly due to complementary charges and the resultant screened electrostatic interaction, can enhance the magnitude and dynamics of protein recruitment to the BL.

### lncRNAs can migrate to the BL to aid recruitment of transcriptional proteins

Since lncRNAs localize at the RL, their concentration profile is peaked at the center of the RL and decays over a length scale of *σ_R_L* = *σ* (Figure S2B). The distance (*L_P_*) between the BL and the RL relative to this length scale is an important regulatory parameter that can affect local lncRNA concentration at the BL and therefore affect protein recruitment (Fig. 3A). Therefore, we wanted to understand how the relative distance (*L_P_/σ*) affects the dynamics of protein recruitment to the BL and condensate formation. It is also important to note that the lncRNA concentration profile can dynamically change due to protein accumulation at the BL, leading to interesting and non-trivial dynamics. We numerically simulated the dynamics described by Eq. 4-5 by varying the distance 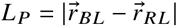 between the loci. We then quantified the protein partitioning to the BL at equilibrium. Protein partitioning to the BL sharply decreases upon increasing the normalized distance *L_P_/σ* (Fig. 3B). When the BL and the RL are in close proximity (small *L_P_/σ*), the protein concentrations at the BL are large enough to form a condensate. At intermediate distances (*L_P_/σ* = 2) which corresponds to the BL and the RL just touching each other, the protein partitioning to the BL begins to decline sharply to a lower value. When the BL and the RL are far away (*L_P_/σ >* 2), the protein partitioning to the BL does not change much and stays at the same low value, which is not enough to form a condensate. In summary, we predict that lncRNAs have a local effect on protein partitioning and condensate formation, that reduces sharply with distance. This local effect is mostly a consequence of lncRNA concentrations decaying over the length scale *σ*, beyond which it has minimal impact on protein recruitment to the BL.

**Figure 3:**
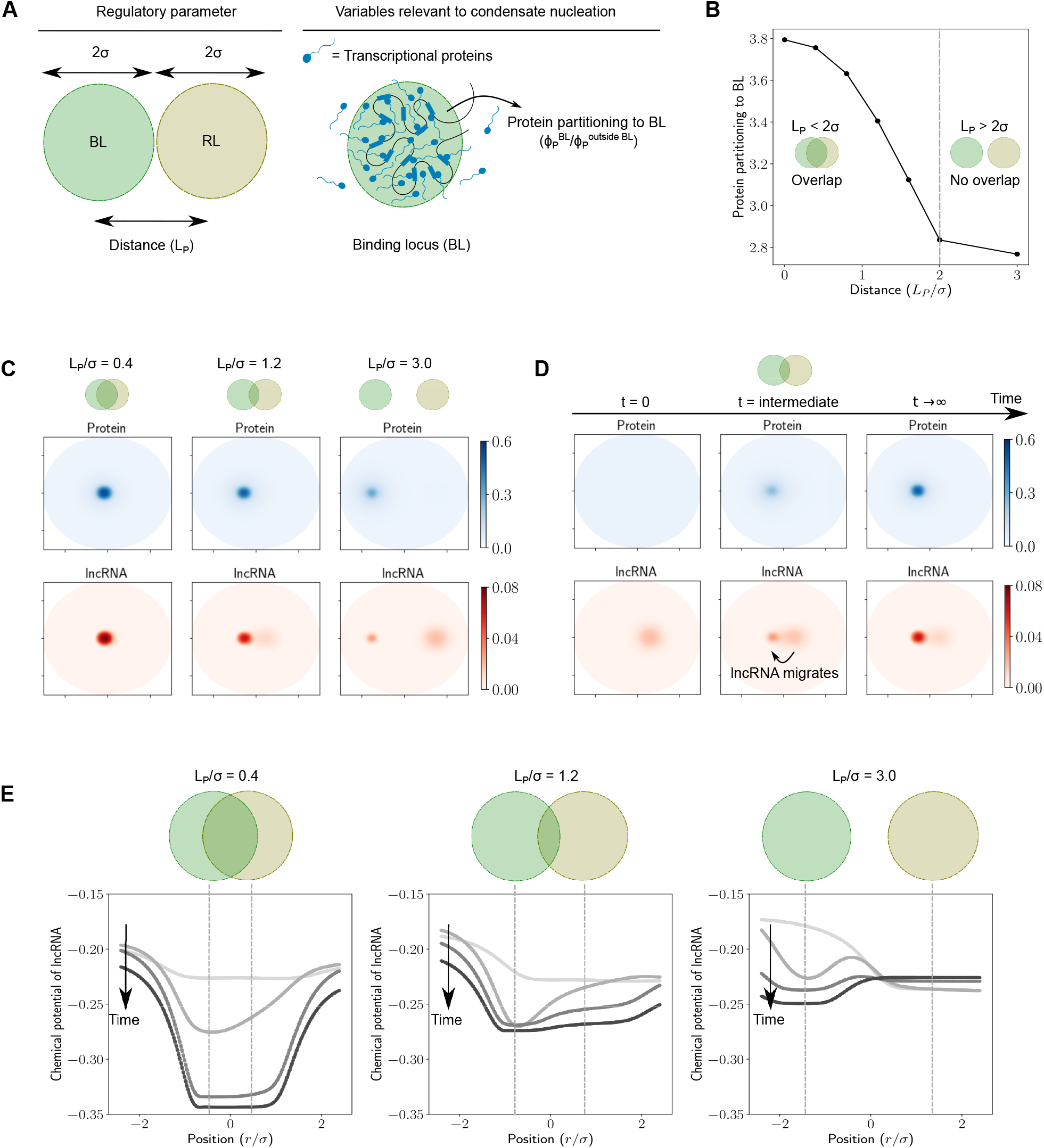
**A** In this figure, we change the distance (*L_P_*) between BL and RL and quantify the protein partitioning to the BL 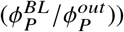. The amount of lncRNA was set to 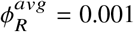 and the average protein concentration to 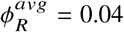 **B** *Condensate formation:* Protein partitioning to the BL upon changing the distance between BL and RL. The distance is reported as the normalized value *L_P_/σ*. When *L_P_/σ* < 2, there is some overlap between the BL and the RL. When *L_P_/σ* > 2, there isn’t any appreciable overlap between the BL and the RL. **C** Concentration profiles of protein and lncRNA at equilibrium for different values of the normalized distance *L_P_/σ* **D** *Dynamics of protein recruitment:* Snapshots of protein and lncRNA concentration profiles at different times. At *t* = 0, the protein is present at a uniform constant concentration everywhere while the lncRNA has a concentration profile peaked at the center of the RL. The distance between the RL and BL is *L_P_/σ* = 1.2, which corresponds to the case with partial overlap. **E** The chemical potential of lncRNA vs. radial position at *t* = 0 for different amounts of lncRNA. The radial position is measured relative to the midpoint of the line connecting the BL and RL, with the origin being the midpoint.

However, this picture is more nuanced when we look at the dynamics. Since the initial lncRNA concentration profile is peaked at the RL and decays with distance, the distance between the BL and the RL affects the initial lncRNA concentration at the BL, and therefore the dynamics of protein recruitment to the BL. To understand this effect, we graphed the concentration profiles of protein and lncRNA for three different values of the scaled distance *L_P_/σ*. At small distances (Fig. 3C, left panel) the RL and the BL are close enough that they almost overlap. The initial lncRNA concentrations at the BL are high because of their proximity to the RL. This helps start a positive feedback cycle, where high lncRNA concentrations at the BL help recruit more protein due to attractive protein-lncRNA interactions, which in turn recruits more lncRNA. This cycle continues until an equilibrium is reached. When the RL and the BL are quite far away (Fig. 3C, right panel), the initial lncRNA concentration at the BL is quite low. In this case, only a small amount of lncRNA migrates from the RL to the BL. Since condensates form only beyond a threshold protein concentration (Fig. 2B), the protein recruited to the BL due to this small amount of lncRNA may not be sufficient to help form a condensate despite the feedback cycle (Fig. 3C). At intermediate distances (Fig. 3C, middle panel) something interesting happens at equilibrium: the lncRNA concentration at the BL seems to be much higher than the RL even though initial lncRNA concentrations at the RL were higher. The time evolution of protein and lncRNA concentration profiles sheds light on this observation (Fig. 3D). At intermediate times, we find that the lncRNA migrates from the RL to the BL. Once this happens, the lncRNA concentration at the BL increases and the positive feedback cycle is initiated, resulting in more protein recruitment.

To understand the mechanistic origin of lncRNA migration, we graphed the chemical potential profile of the lncRNA for different distances (Fig. 3D). This profile dynamically evolves with time. As time progresses, the protein accumulates at the BL because of the attractive well described by Eq. 3. Since proteins attract lncRNAs, increasing protein concentration at the BL makes it an attractive well for the lncRNA which gets deeper with time as proteins accumulate the BL. At short distances (Fig. 3E, left panel), the loci overlap and this well forms essentially at the same location as the RL. Therefore, there is an influx of lncRNA into this region that contains both the RL and the BL. When the distance between the loci is large (Fig. 3E, right panel), not much protein accumulates at the BL initially due to low local lncRNA concentrations. This results in a shallower chemical potential well at the BL for the lncRNA with a chemical potential barrier between the BL and the RL at intermediate times, resulting in a lower migration of lncRNA to the BL. At intermediate distances, there is a partial overlap between the loci (Fig. 3E, middle panel) and the chemical potential for the lncRNA at the BL starts decreasing with protein accumulation at the BL. This leads to a flux of lncRNA away from the RL and into the BL, which is what we see as lncRNA migration.

Given the contrasting effects of the two regulatory parameters - the amount of lncRNA 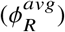 and distance between loci (*L_P_/σ*) - on protein recruitment to the BL, we wanted to understand the impact of them in conjunction (supplemental figure S4). In this figure, the contours correspond to combinations of lncRNA amount and distance that result in the same protein partitioning to the BL. We found that the effect of distance and lncRNA amounts can compensate for each other, resulting in the same value of protein partitioning to the BL for different combinations of these regulatory parameters.

### Non-equilibrium effects result in enhancement or repression of gene expression due to transcription of proximal lncRNAs

The transcription of neighboring lncRNAs can interfere with mRNA transcription by affecting protein concentrations and condensate formation at the BL. Therefore, we next wanted to understand how localized lncRNA transcription from RL affects mRNA transcription from neighboring genes at the BL.

To get a baseline in the absence of lncRNA transcription, we first simulated the model Eq. 6-8 with just mRNA transcription, setting the lncRNA concentrations and transcription rates to zero. We increased the mRNA transcription rate constant *k_M_* and studied the resultant phenomena (section S3 in the supplemental material). Simulations were done using low protein concentrations such that the process of mRNA transcription is necessary for condensate formation. As mRNA is transcribed at the BL, it attracts more protein to the BL, which in turn results in more mRNA transcription since the mRNA transcription rate is coupled to local protein concentration. For a gene expressed at a low level (low *k_M_*), there is not enough mRNA transcription for this positive feedback cycle to recruit enough protein and form a condensate (Fig. S5B, S5C). For moderately expressed genes (moderate *k_M_*), there is enough transcription of mRNA, and the positive feedback cycle results in a stable condensate at steady-state (Fig. S5B, S5C) with a long lifetime (Fig. S5D). For highly expressed genes (large *k_M_*), there is enough mRNA transcription to form a condensate (Fig. S5C). But as mRNA accumulates, the entropic penalty of confining proteins and mRNAs into a dense phase reduces protein concentrations and results in a dissolved condensate at steady-state (Fig. S5B) with a short lifetime (Fig. S5D). These results recapitulate the findings of prior related work in literature (29).

To study the impact of lncRNA transcription on mRNA transcription from the BL, we performed numerical simulations of the model described by Eq. 6-8. The lncRNA transcription rate *k_R_* is an important regulatory parameter here. We increased the lncRNA transcription rate and quantified metrics related to condensate dynamics and gene expression for three different cases: genes expressed at low level i.e. low *k_M_*, genes expressed at moderate level i.e. moderate *k_M_*, and highly expressed genes i.e. high *k_M_* (Fig. 4A).

**Figure 4:**
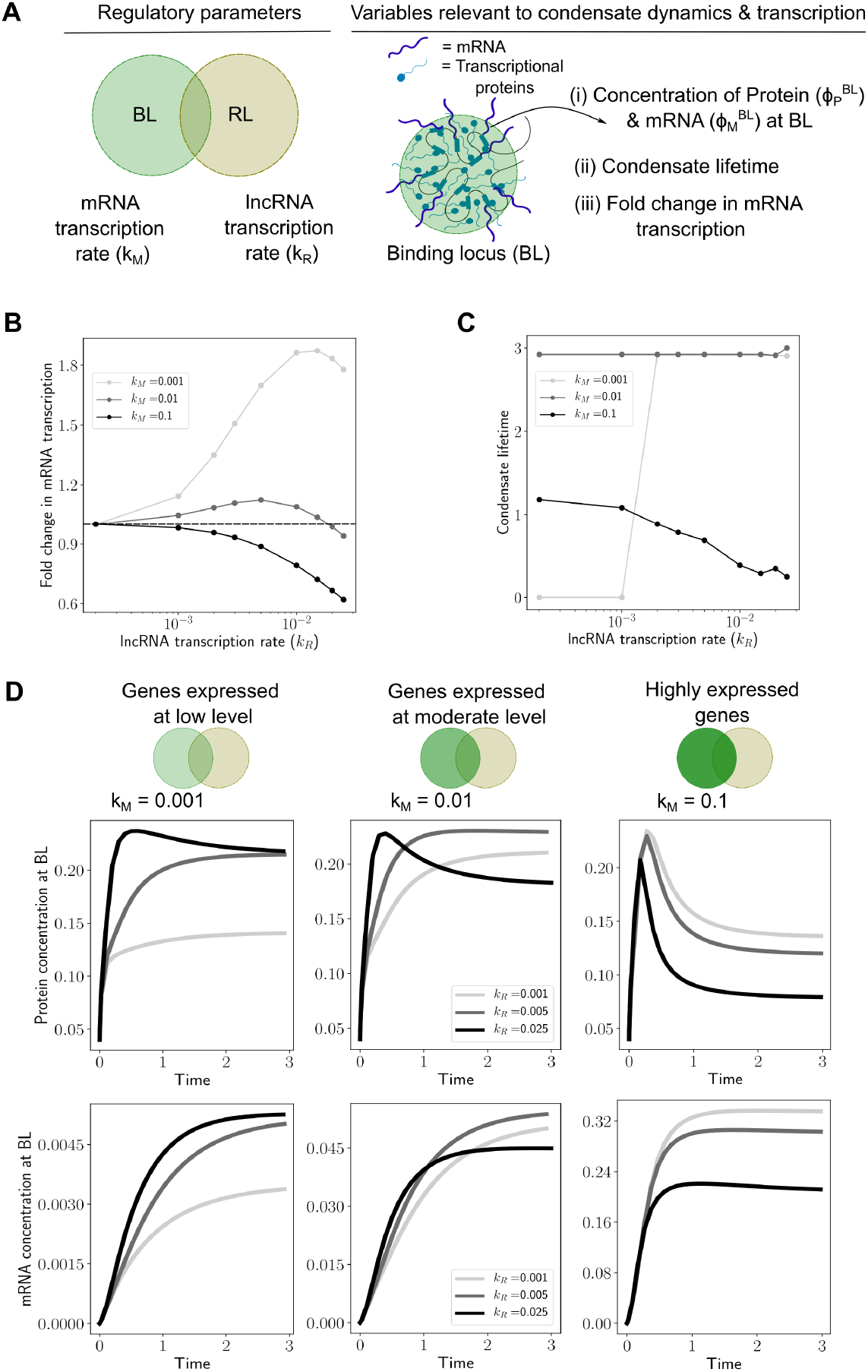
**A** In this figure, we change the transcription rate of the lncRNA (*k_R_*) and study how that impacts condensate dynamics and mRNA transcription for three different regimes of gene expression – (i) genes expressed at low level (*k_M_* = 0.001), (ii) genes expressed at a moderate level (*k_M_* = 0.01), and (iii) highly expressed genes (*k_M_* = 0.1). For each case, we quantified the fold change in mRNA transcription at steady state, condensate lifetime, and the dynamics of protein concentration 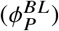 and mRNA concentration 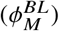 at the BL. For all simulations results in this figure, the distance between the loci was *L_P_/σ* = 0.8, and the protein amount was 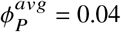 **B Gene expression:** Fold change in mRNA transcription upon changing the lncRNA transcription rate for the three different gene expression regimes. The fold change in mRNA transcription is calculated as = (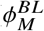 when lncRNA is being transcribed at rate *k_R_*) / (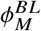 when there is no lncRNA transcription i.e. *k_R_* = 0). The dotted horizontal line corresponds to a fold change value of 1, which means that the lncRNA transcription neither enhances nor represses mRNA transcription. **C Condensate lifetime:** The dependence of condensate lifetime on lncRNA transcription rate for the three different regimes of gene expression. The condensate lifetime is also reported in the dimensionless units (*k_d_t*), and is defined as the duration of time for which protein concentration at the BL is “appreciable”. We chose a cutoff 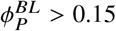 to define “appreciable” protein concentration at the BL. Note that this specific numerical choice of the cutoff value doesn’t change the qualitative nature of the trends or results. **D Dynamics of condensate and gene expression:** Dynamics of protein and mRNA concentration at the BL. Each vertical panel corresponds to a different regime of gene expression. The top panel plots track the protein concentration at the BL with time upon increasing the lncRNA transcription rate (*k_R_*). The bottom panel plots track the mRNA concentration at the BL with time. The time is reported in dimensionless units (*k_d_t*) where *k_d_* is the degradation rate of the mRNA.

For genes expressed at a low level (low *k_M_*), we predict that active transcription of lncRNA at the RL enhances mRNA transcription (Fig. 4B). In this regime, increasing the lncRNA transcription rate leads to an increase in mRNA transcription. This enhancement is accompanied by a corresponding sharp increase in condensate lifetime (Fig. 4C), suggesting that proximal lncRNA transcription enhances protein recruitment to the BL through attractive interactions to form a condensate. This is consistent with the large increase in the protein concentration at the BL at steady state (Fig. 4D, top panel) observed upon increasing the lncRNA transcription rate (*k_R_*) from 0.001 to 0.005. Since the mRNA transcription rate is coupled to protein concentration (Eq. 7), this results in a higher rate of mRNA transcription and therefore higher gene expression, as measured by the steady-state concentration of mRNA (Fig. 4D, bottom panel). However, there are limits to this enhancement in gene expression. Upon further increasing the lncRNA transcription rate *k_R_*, the fold change in mRNA transcription reaches a peak and then reduces (Fig. 4B, *k_M_* = 0.001). This is a consequence of the re-entrant effect of lncRNA concentration on protein condensation. The lncRNA concentration at the BL crosses over from a regime where lncRNA enhances protein recruitment to BL via attractive protein-RNA interactions, to a regime where the lncRNA hinders protein recruitment to the BL due to the entropic costs of confining the proteins and RNAs into a dense phase (Fig. 4B). Transcription of proximal lncRNAs also speeds up response times for gene expression by increasing the initial rate of mRNA transcription (Fig. 4D, bottom panel). The mRNA accumulates more quickly for higher values of *k_R_*, and this is a non-equilibrium effect caused by active lncRNA transcription.

For genes expressed at a moderate level (moderate *k_M_*), active transcription of lncRNA at the RL only has a mild effect on mRNA transcription (Fig. 4B). In this regime, the condensate lifetime is predominantly determined by the dynamics of mRNA transcription and it does not change with increasing *k_R_* (Fig. 4C). The fold change in mRNA transcription has a non-monotonic trend (Fig. 4B). The dynamics of protein and mRNA concentrations at the BL sheds some light on this (Fig. 4D, middle panel). The protein concentration at BL at steady state initially increases and then decreases with *k_R_*. This is again a consequence of switching over to a regime where RNA-RNA repulsion and entropic costs of confining the RNAs and proteins dissolve the condensate. The dynamics (Fig. 4D, middle panel) again reveal that transcription of proximal lncRNAs speeds up response times for gene expression.

For highly expressed genes (moderate *k_M_*), active transcription of lncRNA at the RL has a largely repressive effect on gene expression as the fold change in mRNA transcription monotonically decreases with *k_R_*(Fig. 4B). In this regime, the high *k_M_* already leads to condensate dissolution (Fig. 4D, right panel). lncRNA transcription at the RL further destabilizes condensates as the condensate lifetime decreases with *k_R_* (Fig. 4C). Since increasing *k_R_* reduces the protein concentration at the BL at steady state (Fig. 4D, right panel), this results in slower rates of mRNA transcription and therefore lower gene expression.

In summary, we find that lncRNA transcription has contrasting effects on mRNA transcription from genes expressed at a low level and highly expressed genes. Transcription of proximal lncRNAs increases transcription from the former and represses transcription from the latter. This follows directly from a non-equilibrium model where active lncRNA transcription affects condensate formation at the BL. lncRNA transcription in proximity can alter local RNA concentrations at the BL, which in turn has consequences for protein condensation, and therefore mRNA transcription.

## DISCUSSION

In this study, we propose a simple physical mechanism by which lncRNAs can regulate transcriptional activation and transcription - via attractive interactions with transcriptional proteins that form condensates. Attractive interactions between transcriptional proteins and RNA could arise due to screened electrostatic attraction between oppositely charge polymers (29) which makes this a sequence-agnostic mechanism. At low RNA concentrations, these interactions promote condensation of proteins while high RNA concentrations lead to re-entrant dissolution (Fig. 1B). When coupled with equilibrium mechanisms (e.g. binding) or non-equilibrium mechanisms (e.g. spatially local transcription) that alter their local concentrations, lncRNAs can act as rheostats to fine-tune transcription from neighboring PCGs by regulating transcriptional condensates.

While there has been some experimental work investigating gene regulation by lncRNAs through transcriptional condensates (47), much remains to be understood. Our model makes specific predictions about how different regulatory parameters affect condensate formation, dynamics, and gene expression (Fig. 5), and it serves as a useful conceptual framework to understand many puzzling observations in the literature.

**Figure 5:**
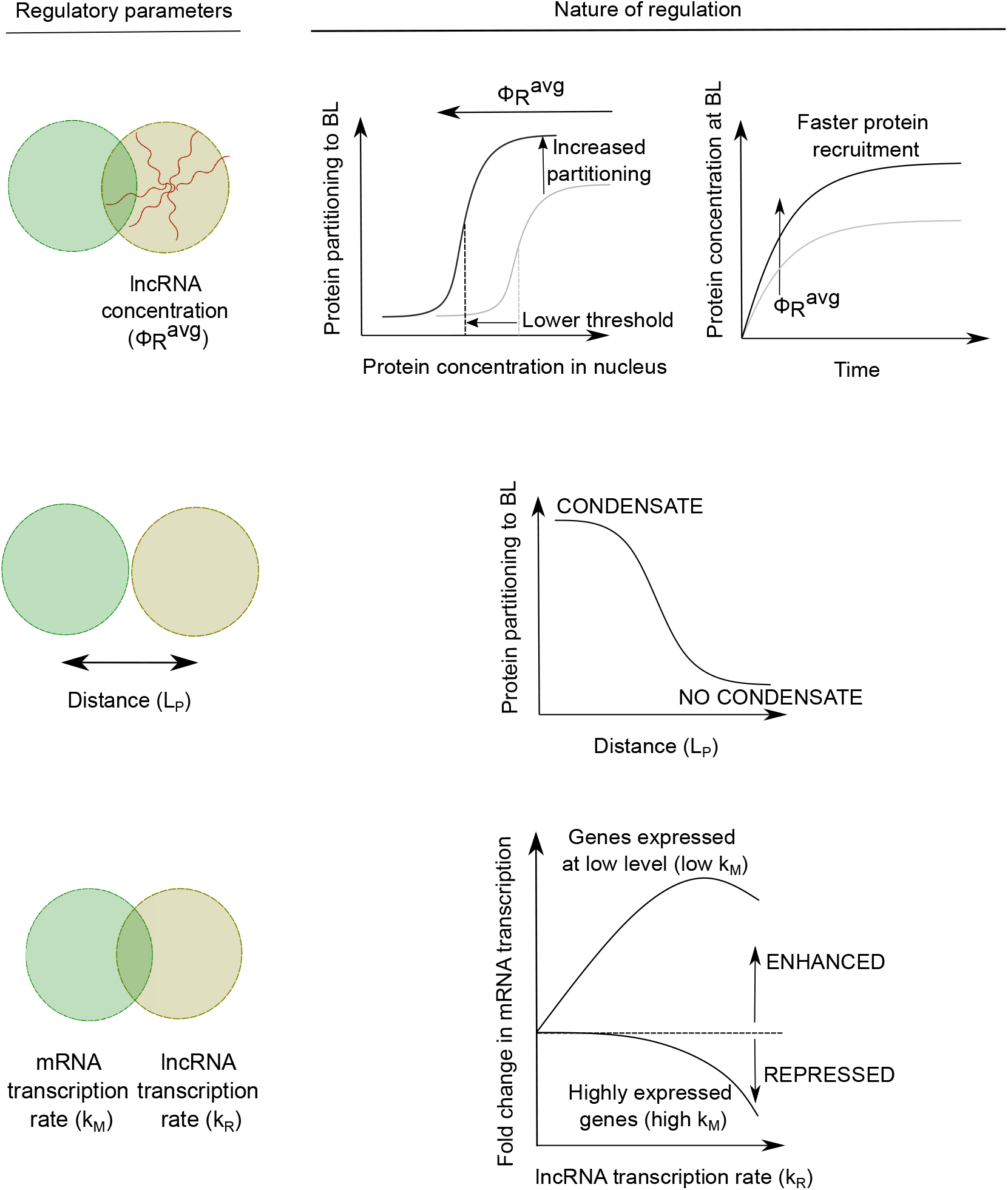
Proximal lncRNAs can regulate condensate formation and mRNA transcription in different ways depending on the regulatory parameter. **A** Increasing the concentration of proximal lncRNAs localized near a BL can bring down the concentration thresholds of transcriptional proteins required for condensate formation, enhance the partitioning of these proteins into the condensate, and speed up protein recruitment to the BL. **B** lncRNAs exert a local effect in enhancing protein partitioning to the BL, and this effect sharply falls off with distance. In some cases, this local effect can be the driving force for condensate formation, with the distance determining whether a condensate will form at the BL or not. **C** Transcription of proximal lncRNAs can increase mRNA transcription from genes expressed at low levels. For highly expressed genes, transcription of proximal lncRNAs represses gene expression.

First, we predict that the presence of a proximal lncRNA near a BL such as a super-enhancer, enhancer, or promoter can reduce threshold protein concentrations required for transcriptional condensate formation, enhance protein partitioning to these loci, and speed up the response time between a stimulus and transcriptional activation.

Second, we predict that the lncRNAs have a spatially local effect on condensate formation, which imposes physical constraints on the spatial and genomic organization of BLs and the lncRNAs that regulate them. This observation can provide a possible explanation for the origin of some known biological facts about lncRNAs. If lncRNAs function by recruiting transcriptional proteins to enhancers and promoters present locally, this can explain why many PCGs are preferentially surrounded by lncRNA-coding loci in their genomic neighborhood (4, 10, 11). Another puzzling fact about lncRNAs is that they have conserved synteny across vertebrates - their genomic positions relative to other genes are conserved rather than their sequence (16). If this local effect of lncRNAs is under evolutionary selection, the effect we predict imposes constraints on the spatial distance between lncRNA-coding genes and promoters. This, together with the observation that syntenic regions in mammals have evolutionarily conserved preferences for spatial contacts (48), can provide a mechanistic explanation for syntenic conservation of lncRNAs across vertebrates (16).

Finally, we predict that proximal transcription of lncRNAs largely represses gene expression from highly transcribed genes while enhancing gene expression from those expressed at a low level. This is also correlated with condensate stability – transcription of proximal lncRNAs enhances gene expression by stabilizing condensates and represses gene expression by destabilizing condensates, depending on the transcription rates of the lncRNA and the mRNA. Experiments that perturb lncRNA amounts and transcription and image condensates and measure gene expression can be used to test this model of whether lncRNAs regulate proximal BLs via interactions with components of transcriptional condensates. This observation also provides a useful framework to understand some conflicting findings in the literature. Studies of transcription regulation by lncRNAs show that they enhance transcription from neighboring PCGs in some cases and inhibit transcription in others (4, 5). Figure 5B gives us a unifying principle that can help reconcile both these observations. For highly expressed genes, transcription of proximal lncRNAs predominantly has a repressive effect as the locally high mRNA concentrations at the BL disfavor condensate formation due to entropic penalties. For genes expressed at low levels, transcription of proximal lncRNAs predominantly enhances gene expression as lncRNAs help attract more protein to the BL via enthalpically favored interactions.

In addition to the regulatory parameters studied in this paper, there is also emerging evidence that RNA secondary structure plays an important role in regulating the formation of biomolecular condensates (49). While we do not explicitly study this effect, our model could be extended to account for this. If we have a description of secondary structures of particular RNA species from experimental techniques such as SHAPE-MaP (50) and if we also know the specific transcriptional proteins this RNA interacts with, we could in principle perform molecular simulations to extract the RNA-protein interaction strength. This can then be an input into our model to perform simulations.

The model itself is agnostic to the identity of RNA species and the principles we identify in this study can be equally applied to understand gene regulation by other kinds of RNA species beyond lncRNAs. For example, this model can be used to understand how an actively transcribing mRNA can lead to transcriptional cross-talk and affect the transcription of neighboring mRNAs. Also, there are several RNA species that can be localized or transcribed near transcriptional condensates including lncRNAs, eRNAs, and divergently transcribed RNAs. These RNAs are often present in low copy numbers in cells (29, 51). Even if the effect of a single locus is mild, several of these RNA loci can act cooperatively to regulate condensate formation and transcription. For example, it is well known that the chromatin is organized into topologically associating domains or TADs (52), which are characterized by a high contact probability of loci within the TAD. Therefore, lncRNAs could cooperatively regulate gene expression within the TAD. Investigating the nature of this cooperative regulation could be an interesting future direction of research.

Our model due to its simplified nature does have several limitations. First, our model is a mean-field description that ignores any stochastic effects that arise due to concentration fluctuations of the protein and RNA species. These fluctuations can be quite important for condensate nucleation and gene expression, and taking them into account can help make additional predictions about how lncRNAs can fine-tune the cell-cell heterogeneity of these phenotypes. Second, our model assumes that the protein concentrations follow Model B dynamics based on a free energy that can be written in terms of the concentration fields. It is quite possible that the dynamics of transcriptional proteins within the dense milieu of condensates with many interacting species can be quite non-trivial and requires other model descriptions. Molecular simulations that model the dynamics of these interacting polymeric species will be required to test whether and when the approximations made in our simplified model break down. Finally, our model assumes that the lncRNA locus and BL do not move much in transcription time scales, which are usually a few minutes for most RNAs. While this is a reasonable approximation given the low diffusivity of the chromatin loci (53), the dynamics of chromatin can couple with the dynamics of transcription and give rise to rich emergent physical phenomena that can provide insights into how transcription shapes genome organization and vice versa. This may be another interesting avenue for future research.

## AUTHOR CONTRIBUTIONS

P.N. and A.K.C conceived the project. P.N. and A.K.C. developed the model. K.S. provided initial code for simulations which was further developed by P.N. P.N. ran numerical simulations and analyzed data. P.N. wrote the first draft of the manuscript and designed the figures. K.S. provided helpful guidance throughout the project. All authors contributed to editing and revising the manuscript.

## ACKNOWLEDGMENTS

P.N. and A.K.C acknowledge support from NSF (Award #2044895). K.S. acknowledges support from the NSF–Simons Center for Mathematical and Statistical Analysis of Biology at Harvard (Award #1764269) and the Harvard Faculty of Arts and Sciences Quantitative Biology Initiative. We thank Jonathan Henninger, Ozgur Oksuz, Kalon Overholt, Richard Young, and Phillip Sharp for several useful discussions.

## SUPPLEMENTARY MATERIAL

An online supplement to this article can be found by visiting BJ Online at http://www.biophysj.org.

