## Supplemental Information for "A model for *cis*-regulation of transcriptional condensates and gene expression by proximal lncRNAs"

### Supporting Material

**S3 Condensate dynamics and mRNA transcription in the absence of actively transcribing lncRNAs** 8

**S4 Supplemental figures** 10

#### S1 Typical distance between lncRNA loci and promoters of BLs/superenhancers

lncRNA-coding loci are usually within a genomic distance of 100 kb from the promoter of protein-coding genes (PCGs) [1]. Superenhancers are typically located at a distance of around 250 kb from the promoters of PCGs [2], which means that lncRNA-coding loci are usually located within a genomic distance of 350 kb of superenhancers. To translate this genomic distance into a spatial distance, we use the worm-like chain model of Beltran et al. [3] for chromatin that takes into account nucleosome heterogeneity and linker DNA. The typical length of linker DNA in euchromatin of human cells is around 50 bp [4], and the Kuhn length of DNA with this linker length is 37.8 nm [3].

Using the size of a nucleotide as 0.34 nm, the number of base pairs present in this Kuhn segment = 37.8 nm/0.34 nm $\times$ (146 bp in nucleosome + 50 bp in linker DNA)/50 bp linker DNA = 436 bp.

Therefore, a 1 kb separation along the genome corresponds to $\approx$ 2 Kuhn segments. Similarly, 350 kb corresponds to $\approx$ 800 Kuhn segments. Using a random polymer model as a crude approximation, (end-end distance = # of Kuhn segments)$^{1/2}$ $\times$ Kuhn length, the average spatial distance between lncRNA-coding loci and promoters/superenhancers comes out to be $\approx 1\ \mu m$. In comparison, super-resolution microscopy studies reveal that transcriptional condensates that form at superenhancers have a diameter in the range of 200-600 nm [5]. Thus, the spatial distance between lncRNA-coding loci and promoters/superenhancers is of the same order as the size of transcriptional condensate.

#### S2 Model description

##### S2.1 Protein-Protein and Protein-DNA interactions

Interactions between intrinsically disordered regions of transcriptional proteins can promote phase separation [6, 7]. These protein-protein interactions that favor phase separation can be qualitatively captured using a mean-field free energy expression from the Flory-Huggins theory of polymer solutions:

$$F_{FH}[\phi_P] = \frac{\phi_P}{N_P}\log\phi_P + (1-\phi_P)\log(1-\phi_P) - \chi_P\phi_P^2 \tag{1}$$

Here, $\phi_P$ is the volume fraction of protein in the solution, $N_P$ is the length of the protein in units of solvent volume, and $\chi_P$ is the Flory-Huggins interaction parameter that captures the magnitude of attractive interactions between protein molecules. Since the volume fraction of the proteins can be converted to protein concentrations by a multiplicative scaling factor, we will be using volume fractions and concentrations as as semantically equivalent while keeping this distinction in mind.

**For this study, we coarse grained the proteins as having $N_P = 5$ beads. To study the phase separation of this protein, the interaction strength $\chi_P$ has to large enough to support phase separation into two phases for some range of concentrations. We chose a value of $\chi_P = 1.1$ for this study.** The corresponding plot of chemical potential with the spinodal and binodal boundaries marked are depicted in figure S1A. Other than this, the particular choice of numerical values of $\chi_P$ and $N_P$ used in this study does not qualitatively affect the phase behavior but only the numerical values of concentration thresholds of the binodal and spinodal boundaries.

In addition to protein-protein interactions, the protein molecules are attracted to regions in space that contain attractive chromatin such as enhancers, super-enhancers and promoters, which we call collectively as the binding loci (BL). These protein-chromatin interactions confer a free energy benefit that is proportional to the protein concentration $\phi_P$ and a Gaussian function of distance from the BL whose $\sigma$ depends on the length scale over which proteins feel the attraction:

$$F_P[\phi_P, \vec{r}] = F_{FH}[\phi_P] + F_{BL}[\phi_P, \vec{r}] = \phi_P \log \phi_P + (1 - \phi_P)\log(1 - \phi_P) - \chi_P \phi_P^2 - c_P e^{-|\vec{r}|^2/\sigma^2} \phi_P \tag{2}$$

The chemical potential associated with this free energy is:

$$\mu_P[\phi_p, \vec{r}] = \frac{\delta F_P}{\delta \phi_P} = \frac{1 + \log \phi_P}{N_P} - (1 + \log(1 - \phi_P)) - 2\chi_P \phi_p - c_P e^{-|\vec{r}|^2/\sigma^2} \tag{3}$$

Figure S1A plots this chemical potential as a function of $\phi_P$ for different values of distance $r$ from the center of the region in space containing DNA binding sites. When the average protein concentration in the system $\phi_P$ is within the spinodal boundary, the free energy $F_P$ is concave function of $\phi_P$ with $\frac{\partial^2 F_P}{\partial \phi_P^2} = \frac{\partial \mu_P}{\phi_P} < 0$. The system is unstable and its free energy can be minimized by splitting into a dense phase rich in protein and a light phase depleted in protein with their respective compositions $\phi_P$ given by the binodal boundary.

S1C and S1D depict the profiles $\phi_p(r)$ in for different values of $\phi_P^{avg}$ and $c_P$. We obtained the equilibrium profiles $\phi_P(r)$ in a circular domain by starting with a spatially uniform protein concentration and simulating the relaxation to equilibrium using Model B dynamics [8]:

$$\frac{\partial \phi_P(\vec{r}, t)}{\partial t} = \vec{\nabla}.\left( D_P \phi_P \left( \vec{\nabla} \frac{\delta F_P}{\delta \phi_P} \right) \right) \tag{4}$$

The steady state solution of the above equations is also the solution to the equation $\mu_p[\phi_P^{eq}, \vec{r}] =$ constant, which is the criterion for chemical equilibrium. The constant is chosen in such a way that $\int \phi_P^{eq}(\vec{r}) dV = \phi_P^{avg}$. Expanding out the equation:

$$\mu_P[\phi_P^{eq}, \vec{r}] = \frac{1 + \log \phi_P^{eq}}{N_P} - (1 + \log(1 - \phi_P^{eq})) - 2\chi_P \phi_P^{eq} - c_P e^{-|\vec{r}|^2/\sigma^2} = \text{constant} \tag{5}$$

Consider the system having average protein concentration $\phi_P^{avg}$ well outside the spinodal boundary. Assuming a large system, $\phi_P^{avg}$ is going to be the protein concentration at $r \to \infty$. The spatially varying free energy benefit $F_{BL}[\phi_P, \vec{r}]$ conferred by the protein-DNA interactions leads to an accumulation of protein at the center $r = 0$ of the spatial region containing DNA binding sites. If $\phi_P^{avg}$ or $c_P$ is sufficiently high, then $\phi_P^{eq}(r = 0)$ can cross the spinodal boundary and a dense phase of protein starts to form at $r = 0$. In figure S1C and S1D, we observe that a small increase in $\phi_P^{avg}$ from 0.08 to 0.1 and a small increase in $c_P$ from 0.05 to 0.1 results in a large increase in $\phi_P^{eq}(r = 0)$. For a given $\phi_P^{avg}$, sufficiently strong protein-DNA interactions with $c_P > c_P^*(\phi_P^{avg})$ can result in the formation of a dense phase of protein at much lower protein concentrations well below the spinodal boundary. We interpret the dense phase of protein to represent a transcriptional condensate.

Figure S1B plots contours of condensate area, which is the area of a spatial region with $\phi_P^{eq}(\vec{r}) > 0.3$, as we vary $\phi_P^{avg}$ and $c_P$ in the simulations. When we have a sufficient amount of protein $\phi_P^{avg}$ in the system or sufficiently strong magnitude $c_P$ of the protein-DNA interactions, a condensate of non-zero area is nucleated. The white dotted line represents the curve $c_P = c_P^*(\phi_P^{avg})$. The simulations agree with our theoretical prediction that a condensate nucleates when $c_P > c_P^*(\phi_P^{avg})$.

**Simulations for results in the main manuscript are done in a regime close to the critical boundary $c_P = c_P^*(\phi_P^{avg})$ to illustrate how the presence of other RNA species can alter the condensate formation process. Therefore, simulations results in the main manuscript are done for the parameters $c_p = 0.2, \phi_P^{avg} = 0.04$ unless stated otherwise.**

##### S2.2 RNA-RNA and RNA-DNA interactions

Recent studies show that long-non coding RNAS (lncRNAs) tend to be present in locally high concentrations near the DNA loci (RL) that code for them [9]. Although the mechanisms that cause this localization is poorly understood, it could be a consequence of:

1. Equilibrium effects such as tethering of lncRNAs to their DNA loci by proteins such as Polymerase II [10] or YY1 [11]
2. Non-equilibrium effects such as localized production of lncRNAs coupled with diffusion that can result in higher concentration of lncRNAs near the regions of lncRNA production

In this section, we will model equilibrium mechanisms that keep lncRNAs bound to their DNA coding loci using similar arguments as section S2.1. The free energy of binding of lncRNAs to their DNA loci is:

$$F_{RL}(\phi_R, \vec{r}) = -c_R e^{-|\vec{r}|^2/\sigma^2} \phi_R \tag{6}$$

where $\phi_R$ is the concentration of RNA and $c_R$ is a parameter that captures the strength of the RNA-DNA interactions. In addition, RNA-RNA repulsion and entropy of the RNA polymer and solvent also contribute to the total free energy of a system containing RNA, its DNA locus, and solvent:

$$F_R[\phi_R, \vec{r}] = \frac{\phi_R}{N_R} \log \phi_R + (1 - \phi_R) \log(1 - \phi_R) + \chi_R \phi_R^2 - c_R e^{-|\vec{r}|^2/\sigma^2} \phi_R \tag{7}$$

Here, $N_R$ is the length of the RNA polymer and $\chi_R$ is the magnitude of the repulsive strength between the RNA species. **The typical length of disordered regions of transcriptional proteins are not more than $1000$ amino acids [12]. On the other hand, lncRNAs and mRNAs have a length of the order of magnitude $\sim 10000$ base pairs, around 10 times longer [13]. Therefore, we set the length of the RNA polymer as $N_R = 50$ beads. The strength of the repulsive interactions with the RNA species was set to a value of $\chi_R = 2.0$, which is stronger than the protein-protein attractive interactions. The strength of the RNA-DNA interactions was set to a similar value as the protein as $c_R = 0.2$. This actually represents an upper limit on the strength of lncRNA-DNA interactions mediated by chromatin-binding proteins. Transcription factors bind to the DNA with a binding affinity in the 1-100 nM range [14] while the binding affinity of lncRNAs with chromatin binding proteins is slightly weaker, around 100-1000 nM [15]. Assuming a similar density of binding sites for lncRNA and tethering proteins on the DNA, $c_R$ is typically much lower than $c_P$.** The chemical potential associated with this free energy is:

$$\mu_R[\phi_R, \vec{r}] = \frac{\delta F_R}{\delta \phi_R} = \frac{1 + \log \phi_R}{N_R} - (1 + \log(1 - \phi_R)) + 2\chi_R \phi_R - c_R e^{-|\vec{r}|^2/\sigma^2} \tag{8}$$

Figure S2A plots this chemical potential as a function of $\phi_R$ for different values of distance $r$ from the center of the region in space containing DNA binding sites. Figure S2B depicts the equilibrium profiles $\phi_R(r)$ in for different values of $\phi_R^{avg}$. We obtained the equilibrium profiles $\phi_R(r)$ in a circular domain by starting with a spatially uniform RNA concentrations $\phi_R^{avg}$ and simulating the relaxation to equilibrium using Model B dynamics [8]:

$$\frac{\partial \phi_R(\vec{r})}{\partial t} = \vec{\nabla}.\left( D_R \phi_R \left( \vec{\nabla} \frac{\delta F_R}{\delta \phi_R} \right) \right) \tag{9}$$

The steady state solution of the above equations is also the solution to the equation $\mu_R[\phi_R^{eq}, \vec{r}] =$ constant, which is the criterion for chemical equilibrium. The constant is chosen in such a way that $\int \phi_R^{eq}(\vec{r}) dV = \phi_R^{avg}$. Expanding out the equation:

$$\mu_R[\phi_R^{eq},\vec{r}] = \frac{1+\log\phi_R^{eq}}{N_R} - (1+\log(1-\phi_R^{eq})) + 2\chi_R\phi_R^{eq} - c_R e^{-|\vec{r}|^2/\sigma^2} = \text{constant} \tag{10}$$

For low values of $\phi_R$, the dominant $\phi_R$-dependent term in the chemical potential is $\frac{\log\phi_R}{N_R}$, which comes from the entropy of RNA in solution. The equilibrium concentration profile is shaped by the balance between RNA-DNA interactions that attract the RNA to its DNA locus and thermal fluctuations that tend to equalize concentrations everywhere. In this regime, the equilibrium profile $\phi_R^{eq}(r)$ is obtained by solving the equation:

$$\mu_R[\phi_R^{eq},\vec{r}] \approx \frac{1+\log\phi_R^{eq}}{N_R} - c_R e^{-|\vec{r}|^2/\sigma^2} = \text{constant} \tag{11}$$

For intermediate values of $\phi_R$, the dominant $\phi_R$-dependent term in the chemical potential is $2\chi_R\phi_R$, which comes from the RNA-RNA repulsive interactions. The equilibrium concentration profile is shaped by the balance between RNA-DNA interactions that attract the RNA to its DNA locus and RNA-RNA repulsions that tend to equalize concentrations everywhere. In this regime, the equilibrium profile $\phi_R^{eq}(r)$ is obtained by solving the equation:

$$\mu_R[\phi_R^{eq},\vec{r}] \approx 2\chi_R\phi_R^{eq} - c_R e^{-|\vec{r}|^2/\sigma^2} = \text{constant} \tag{12}$$

In this regime, the free energy penalty imposed by RNA-RNA repulsion linearly scales with $\phi_R$. Therefore, the DNA locus get saturated with a fixed amount of RNA and any additional RNA added will get uniformly distributed across the system. This is the reason that the profiles $\phi_R^{eq}(r)$ in figure S1F for intermediate values of $\phi_R^{avg} = 0.005$ and $\phi_R^{avg} = 0.01$ maintain their Gaussian shape while just being shifted up by some constant amount.

##### S2.3 Protein-RNA interactions

The disordered regions of many transcriptional proteins contain a net positive charge. They can attract negatively charged RNAs via screened electrostatic interactions [12]. Prior studies have shown that the qualitative features of phase diagrams of charged polymers in solutions interacting via screened electrostatic interactions can be qualitatively captured via a mean-field Flory Huggins free energy expression [16]. For a Protein-RNA solution, the free energy expression can be written down to be:

$$F_{FH}[\phi_P,\phi_R] = \frac{\phi_P}{N_P}\log\phi_P + \frac{\phi_R}{N_R}\log\phi_R + (1-\phi_P-\phi_R)\log(1-\phi_P-\phi_R) - \chi_P\phi_P^2 - \chi_{PR}\phi_P\phi_R + \chi_R\phi_R^2 \tag{13}$$

Here, $\chi_P$ captures the magnitude of protein-protein attractive interactions, $\chi_{PR}$ captures the magnitude of the RNA-Protein screened electrostatic attraction and $\chi_R$ captures the magnitude of the screened RNA-RNA electrostatic repulsion. We already talked about the numerical values of the parameters $\chi_P$ and $\chi_R$ in the previous sections. **We need a sufficiently large value of the attraction strength $\chi_{PR}$ that leads to an increase in protein partitioning to the dense phase when we have some RNA in the system (i.e. $\phi_R > 0$), but not too large enough to preclude the dissolution of the dense phase of protein, as this is the observed biological phenomenology [12]. The value $\chi_{PR} = 1.2$ was used for these simulations, and it satisfies this phenomenology as shown in figure S3C.**

The solvent entropy term $(1-\phi_P-\phi_R)\log(1-\phi_P-\phi_R)$ can be rearranged as:

$$\begin{aligned}(1-\phi_P-\phi_R)\log(1-\phi_P-\phi_R) &= (1-\phi_P-\phi_R)\Big[\log(1-\phi_P) + \log\Big(1-\frac{\phi_R}{1-\phi_P}\Big)\Big] \\ &= (1-\phi_P)\log(1-\phi_P) + (1-\phi_P)\log\Big(1-\frac{\phi_R}{1-\phi_P}\Big) - \phi_R\log(1-\phi_P) - \phi_R\log\Big(1-\frac{\phi_R}{1-\phi_P}\Big)\end{aligned} \tag{14}$$

At dilute RNA and Protein concentrations $\phi_P << 1$ and $\phi_R << 1$, using the expansions $\log(1-x) = -x - x^2/2 + ...$ and $1/(1-x) = 1 + x + x^2 + ...$, the above terms can be expanded to yield the following expression for solvent entropy:

$$(1-\phi_P-\phi_R)\log(1-\phi_P-\phi_R) = (1-\phi_P)\log(1-\phi_P) + \phi_R\phi_P + \frac{\phi_R^2}{2} + \frac{\phi_R^2\phi_P}{2} + \frac{\phi_R\phi_P^2}{2} + \frac{\phi_R^2\phi_P^2}{2} + ... \tag{15}$$

Under this approximation, the free energy in equation 13 gets modified as:

$$F_{FH,mod}[\phi_P, \phi_R] = \frac{\phi_P}{N_P}\log\phi_P + (1-\phi_P)\log(1-\phi_P) - \chi_P\phi_P^2$$
$$+ \frac{\phi_R}{N_R}\log\phi_R + (1-\chi_{PR})\phi_P\phi_R + \Big(\chi_R + \frac{1}{2}\Big)\phi_R^2 + \frac{\phi_R^2\phi_P + \phi_R\phi_P^2 + \phi_R^2\phi_P^2}{2} \quad (16)$$

The above free energy expression has nicer numerical properties and leads to lower numerical errors compared to the free energy expression 13 when simulating the dynamical equations in sections S2.5 and S2.7. Therefore, we will use this free energy expression for the rest of the study.

But before doing this, let us first establish that both $F_{FH}$ and $F_{FH,mod}$ do not lead to any qualitative difference in the equilibrium RNA-protein phase diagram due to the approximations introduced. From the free energy expressions 13 and 16, we can generate the phase diagram by analyzing the Jacobian matrix of $F_{FH}[\phi_P, \phi_R]$ and $F_{FH,mod}[\phi_P, \phi_R]$ and with respect to the variables $\phi_P$ and $\phi_R$:

$$J_{FH} = \begin{bmatrix} \frac{\partial^2 F_{FH}}{\partial\phi_P^2} & \frac{\partial^2 F_{FH}}{\partial\phi_P\partial\phi_R} \\ \frac{\partial^2 F_{FH}}{\partial\phi_P\partial\phi_R} & \frac{\partial^2 F_{FH}}{\partial\phi_R^2} \end{bmatrix} = \begin{bmatrix} \frac{1}{N_P\phi_P} + \frac{1}{1-\phi_P-\phi_R} - 2\chi_P & \frac{1}{1-\phi_P-\phi_R} - \chi_{PR} \\ \frac{1}{1-\phi_P-\phi_R} - \chi_{PR} & \frac{1}{N_R\phi_R} + \frac{1}{1-\phi_P-\phi_R} + 2\chi_R \end{bmatrix} \quad (17)$$

$$J_{FH,mod} = \begin{bmatrix} \frac{\partial^2 F_{FH,mod}}{\partial\phi_P^2} & \frac{\partial^2 F_{FH,mod}}{\partial\phi_P\partial\phi_R} \\ \frac{\partial^2 F_{FH,mod}}{\partial\phi_P\partial\phi_R} & \frac{\partial^2 F_{FH,mod}}{\partial\phi_R^2} \end{bmatrix} = \begin{bmatrix} \frac{1}{N_P\phi_P} + \frac{1}{1-\phi_P} - 2\chi_P + \phi_R + \phi_R^2 & 1 - \chi_{PR} + \phi_R + \phi_R + 2\phi_P\phi_R \\ 1 - \chi_{PR} + \phi_R + \phi_R + 2\phi_P\phi_R & \frac{1}{N_R\phi_R} + (2\chi_R + 1) + \phi_P + \phi_P^2 \end{bmatrix} \quad (18)$$

The system is unstable and can undergo phase separation in the regions in the $\phi_P - \phi_R$ space where the Jacobian has at least one negative eigenvalue, which corresponds to a region of concavity of the free energy. The brown region in figure S3F corresponds to this region of spinodal instability. When $\phi_P$ and $\phi_R$ are within this region, the system splits into two phases: a dense phase rich in RNA and protein and a light phase poor in both RNA and protein. The coexistence compositions $(\phi_P^{dense}, \phi_R^{dense})$ and $(\phi_P^{light}, \phi_R^{light})$ are obtained by solving the equations for equality of chemical potentials and osmotic pressures in the two phases, which form the criteria for multiphase equilibrium:

$$\mu_P(\phi_P^{dense}, \phi_R^{dense}) = \mu_P(\phi_P^{light}, \phi_R^{light}) \quad (19)$$
$$\mu_R(\phi_P^{dense}, \phi_R^{dense}) = \mu_R(\phi_P^{light}, \phi_R^{light}) \quad (20)$$
$$\Pi(\phi_P^{dense}, \phi_R^{dense}) = \Pi(\phi_P^{light}, \phi_R^{light}) \quad (21)$$

where the chemical potentials of protein and RNA are respectively $\mu_P = \frac{\partial f_{FH}}{\partial\phi_P}$, $\mu_R = \frac{\partial f_{FH}}{\partial\phi_R}$, and the osmotic pressure $\Pi = k_B T/V_{solution}\Big(f_{FH} - \mu_P\phi_P - \mu_R\phi_R\Big)$. In addition, the total amount of protein ($\phi_P^{avg}$) and RNA ($\phi_R^{avg}$) constrain the dense and light phase compositions in the following way:

$$\nu\phi_P^{dense} + (1-\nu)\phi_P^{light} = \phi_P^{avg} \quad (22)$$
$$\nu\phi_R^{dense} + (1-\nu)\phi_R^{light} = \phi_R^{avg} \quad (23)$$

The system of 5 equations 19- 23, need to be solved to get the variables $\phi_P^{dense}$, $\phi_P^{light}$, $\phi_R^{dense}$, $\phi_R^{light}$, and $\nu$. Here, $\nu$ is the volume fraction of the dense phase.

Figures S3A and S3B show the phase diagram for the two different free energy expressions 13 and 16 using the procedure described above. For different starting compositions in the spinodal region, the red dotted lines represent the coexistence curves with their ends being the dense phase and the light phase compositions. The positive slope of these dotted lines indicates that the dense phase is rich in both RNA and protein, while the light phase is depleted in both. We can see that the shapes of the binodal and spinodal boundaries and the slopes of the tie-lines connecting the coexisting concentrations are qualitatively the same. Therefore, we will use $F_{FH,mod}$ for the rest of the study.

Beyond a critical value i.e. $\phi_R > \phi_R^c = 0.17$, there is no phase separation and formation of a two-phase region. As we titrate $\phi_R$ in the system for a constant $\phi_P = 0.3$, we observe that the horizontal width of the coexistence curve initially increases and then decreases. This phenomenon is illustrated more clearly in figure S3C where we compute the protein partition ratio $\phi_P^{dense}/\phi_P^{light}$, which also shows the same trend upon titrating the system with $\phi_R$. RNA promotes the partitioning of the protein into the dense phase at low $\phi_R$ by virtue of its attractive interactions with the protein. At high $\phi_R$, the RNA-RNA repulsive interactions and the entropic penalty of excluding the solvent from the dense phase results makes the partitioning of protein into a dense phase unfavorable.

In this way, a simple Flory-Huggins model captures the qualitative features of a re-entrant phase diagram arising due to RNA-Protein screened electrostatic interactions that have been observed in other studies [17, 18, 12, 19, 20].

##### S2.4 Parameters associated with the free energy

| Parameter | Value | Description | Rationalization |
|---|---|---|---|
| $N_P$ | 5.0 | Length of coarse-grained protein sequence | Section S2.1 |
| $c_P$ | 0.2 | Protein-BL interaction strength | Section S2.1 |
| $\chi_P$ | 1.1 | Protein-protein attraction strength | Section S2.1 |
| $N_R$ | 5.0 | Length of coarse-grained lncRNA sequence | Section S2.2 |
| $c_R$ | 0.2 | lncRNA-RL interaction strength | Section S2.2 |
| $N_M$ | 5.0 | Length of coarse-grained mRNA sequence | Coarse-grained mRNA and lncRNA sequences have lengths of order 10 kb [13] |
| $\chi_R$ | 2.0 | RNA-RNA repulsion strength | Section S2.1 |
| $\chi_{PR}$ | 1.2 | RNA-protein interaction strength | Section S2.3 |
| $\phi_P^c$ | 0.15 | Protein concentration threshold to determine dense phase/condensate | Figure S3B |
| $\kappa$ | 0.1 | Surface tension | Set to a small value that ensures smooth protein phase boundaries |

Table S1: Table of parameters associated with the free energy expression

A value of $\sigma_{BL} = 5$ was chose by trial and error to ensure that the diameter of the condensate to the diameter of the simulation domain, which was set to 30 units, had approximately the same ratio as the diameter of a transcriptional condensate ($\sim 0.6$ nm) [5] to the diameter of a eukaryotic nucleus ($\sim 8\mu m$). For simplicity, we chose $\sigma_{RL} = \sigma_{BL} = \sigma = 5$. In principle, this can be varied and studied in more detail.

##### S2.5 Dynamics of condensate formation

We argued earlier in section S2 that the diffusion of transcriptional proteins happens over a faster time scale of $\tau_P \sim 1s$. At these time scales, RNAs are not being turned over and the total amount of RNAs in the system can be considered a conserved parameter. The transcriptional proteins present at a uniform concentration within the nucleus are activated and gain the ability to bind to the chromatin. In addition, they can also interact with the RNAs through the screened electrostatic interactions discussed above. Taking into account all of these interactions, the overall free energy of this system is:

$$F[\phi_P,\phi_R] = \underbrace{\frac{\phi_P}{N_P}\log\phi_P + \frac{\phi_R}{N_R}\log\phi_R + (1-\phi_P-\phi_R)\log(1-\phi_P-\phi_R)}_{\text{Entropy}} - \underbrace{\chi_P\phi_P^2}_{\text{Protein-Protein}} - \underbrace{\chi_{PR}\phi_P\phi_R}_{\text{Protein-RNA}} + \underbrace{\chi_R\phi_R^2}_{\text{RNA-RNA}}$$
$$- \underbrace{c_P e^{-|\vec{r}-\vec{r}_{BL}|^2/\sigma^2}\phi_P}_{\text{Protein-chromatin}} - \underbrace{c_R e^{-|\vec{r}-\vec{r}_{RL}|^2/\sigma^2}\phi_R}_{\text{RNA-DNA}} + \underbrace{\frac{\kappa}{2}|\nabla\phi_P|^2}_{\text{Surface Tension}} \quad (24)$$

Using the approximation for the solvent entropy (equation 15), the free energy can be rewritten as:

$$F[\phi_P,\phi_R] = \frac{\phi_P}{N_P}\log\phi_P + (1-\phi_P)\log(1-\phi_P) - \chi_P\phi_P^2 + \frac{\phi_R}{N_R}\log\phi_R + (1-\chi_{PR})\phi_P\phi_R$$
$$+ \left(\chi_R + \frac{1}{2}\right)\phi_R^2 + \frac{\phi_R^2\phi_P + \phi_R\phi_P^2 + \phi_R^2\phi_P^2}{2} - c_P e^{-|\vec{r}-\vec{r}_{BL}|^2/\sigma^2}\phi_P - c_R e^{-|\vec{r}-\vec{r}_{RL}|^2/\sigma^2}\phi_R + \frac{\kappa}{2}|\nabla\phi_P|^2 \quad (25)$$

The field $\phi_P(\vec{r})$ relaxes back to a new equilibrium state as a consequence of these interactions. The RNA species that are localized nearby due to its interactions with DNA also diffuse and reorganize themselves in response to temporal changes in $\phi_P$. The coupled dynamics of relaxation to equilibrium can be captured using the following Model B equations [8]:

$$\frac{\partial \phi_P(\vec{r})}{\partial t} = \vec{\nabla}.\left( D_P \phi_P \Big( \vec{\nabla} \frac{\delta F}{\delta \phi_P} \Big) \right) \tag{26}$$

$$\frac{\partial \phi_R(\vec{r})}{\partial t} = \vec{\nabla}.\left( D_R \phi_R \Big( \vec{\nabla} \frac{\delta F}{\delta \phi_R} \Big) \right) \tag{27}$$

While the equilibrium profiles $\phi_P^{eq}(\vec{r})$ and $\phi_R^{eq}(\vec{r})$ can be obtained by solving $\mu_P[\phi_P^{eq}, \phi_R^{eq}, \vec{r}]$ = constant and $\mu_R[\phi_P^{eq}, \phi_R^{eq}, \vec{r}]$ = constant.

##### S2.6 Parameters associated with dynamics

In the model equations $D_P, D_R$ and $D_M$ are the diffusivity of the transcriptional proteins, lncRNAs, and mRNAs respectively in dilute solution. For lncRNAs and mRNAs that are being actively transcribed, we assume that they are strongly tethered to the chromatin by RNA Polymerase II, and their diffusivity is the same as diffusivity of the BL or the RL [12]. Diffusivity of actively transcribed chromatin loci are of the order $10^{-2} \mu m^2/s$ [21] which is about 1000 times smaller than the diffusivity of transcriptional proteins. Therefore, we set $D_P = 100$ and $D_R = D_M = 0.1$ for our simulations.

$k_{dR}$ and $k_{dM}$ are the first-order degradation rates of the lncRNA and the protein respectively. The half-lives of RNAs span a range of time scales from minutes to hours. However, the median half-lives are not that different and are of the same order of magnitude for both mRNAs and lncRNAs – both being a few hours [22]. Therefore, we set both degradation rate constants to the same value $k_{dM} = k_{dR} = k_d = 0.02$. This value was chosen such that the half-life of the RNA species $\ln 2/k_d \approx 35$ is an order of magnitude larger than the protein diffusion time scale $\tau_D = r^2/D_P = 2.25$. This is consistent with biological reality where proteins diffuse much faster than the RNA half-lives.

The parameters $k_M$ and $k_R$ quantify the magnitude of mRNA and lncRNA transcription rates and $\sigma_M$ and $\sigma_R$ refer to the spatial extent of these molecules. Since the mean lengths of lncRNAs and mRNAs in the human genome are of the same order of magnitude [23] – around 10 kb, we set $\sigma_M = \sigma_R = \sigma = 5$. The value of $k_M$ and $\sigma$ dictates the condensate morphology. Intermediate values of $k_M$ and $\sigma$ favor condensate formation and spherical morphologies. When $k_M$ is high, we get vacuolar morphologies and aspherical morphologies when $\sigma$ is large and condensate dissolution when $\sigma$ is small [24].

| Parameter | Value | Description |
|---|---|---|
| $D_P$ | 100 | Protein diffusivity |
| $D_R$ | 0.01 | lncRNA diffusivity |
| $D_M$ | 0.01 | mRNA diffusivity |
| $\sigma_R$ | 5 | Spatial extent of lncRNA locus |
| $\sigma_M$ | 5 | Spatial extent of mRNA locus |
| $k_{dR}$ | 0.02 | lncRNA degradation rate |
| $k_{dM}$ | 0.02 | mRNA degradation rate |
| ‘ | | |

Table S2: Table of parameters associated with dynamical equations

##### S2.7 Dynamics of active transcription

Active transcription and depletion of RNAs can change the RNA concentrations and provide a driving force that pushes the system out of equilibrium. In our model, we make a distinction between two kinds of RNAs - (i) mRNAs which are transcribed starting from the promoter of protein-coding genes in spatial regions where transcriptional condensates form. The production rates of these are coupled to the local protein concentrations $\phi_P$ and (ii) lncRNAS, which are transcribed from nearby DNA present in the vicinity of BL. The production rates of these RNAs in general do not depend on the transcriptional proteins that form at BLs to activate transcription and their transcription rate is independent of $\phi_P$.

To first understand the effect of localized mRNA transcription on the dynamics of transcriptional condensates, we model the dynamics of $\phi_P(\vec{r},t)$ using Model B dynamics [8] and couple this to a reaction-diffusion model for the dynamics of the concentration field that corresponds to mRNA ($\phi_M$):

$$\frac{\partial \phi_P(\vec{r},t)}{\partial t} = \vec{\nabla}.\Bigg(D_P\phi_P\Big(\vec{\nabla}\frac{\delta F}{\delta \phi_P}\Big)\Bigg) \tag{28}$$

$$\frac{\partial \phi_M(\vec{r},t)}{\partial t} = D_M\nabla^2\phi_M + k_M e^{\frac{-|\vec{r}-\vec{r}_{BL}|^2}{\sigma^2}}\phi_P - k_d\phi_M \tag{29}$$

From the above equations: (i) the dynamics of the field $\phi_P(\vec{r},t)$ is coupled to the dynamics of the field $\phi_M(\vec{r},t)$ via the protein-RNA interactions captured by the free energy $F$ (ii) the dynamics of the field $\phi_M(\vec{r},t)$ is coupled to the dynamics of the field $\phi_P(\vec{r},t)$ as the rate of production of the RNA depends on $\phi_P(\vec{r},t)$. These couplings result in RNA production acting as a feedback on the protein transport, resulting in different non-equilibrium steady states depending on the system parameters.

Next, we would like to understand how the transcriptional dynamics of lncRNAs produced near transcriptional condensates interferes and affects the dynamics of $\phi_P(\vec{r},t)$. To study this, we compare the dynamics described by 28, 29 with the below equations in the presence of the second RNA species i.e. lncRNAs, by progressively increasing the lncRNA transcription rate:

$$\frac{\partial \phi_P(\vec{r},t)}{\partial t} = \vec{\nabla}.\Bigg(D_P\phi_P\Big(\vec{\nabla}\frac{\delta F}{\delta \phi_P}\Big)\Bigg) \tag{30}$$

$$\frac{\partial \phi_M(\vec{r},t)}{\partial t} = D_M\nabla^2\phi_M + k_M e^{\frac{-|\vec{r}-\vec{r}_{E}|^2}{\sigma^2}}\phi_P - k_d\phi_M \tag{31}$$

$$\frac{\partial \phi_R(\vec{r},t)}{\partial t} = D_R\nabla^2\phi_R + k_R e^{\frac{-|\vec{r}-\vec{r}_{R}|^2}{\sigma^2}} - k_d\phi_R \tag{32}$$

Here, the rate constants $k_M(\vec{r})$ and $k_R(\vec{r})$ are modeled as spatially dependent Gaussians centered around the BL locus and the lncRNA locus. The peak values of these Gaussians are $k_M$ and $k_R$ and their widths are both fixed to be the same value $\sigma$ for simplicity.

In the above model, the transcription of the proximal lncRNAs can perturb the the dynamics of the base system described by equations 28, 29. The extent of this perturbation will depend on the parameters $k_M$ which couples the dynamics of the field $\phi_M(\vec{r},t)$ to the field $\phi_P(\vec{r},t)$.

##### S2.8 Formulae to calculate different quantities to analyze simulation results

In the below expressions, the species $s$ could refer to transcriptional proteins, lncRNA, or mRNA.

- Concentration of a species $s$ at the BL = $\phi_s^{BL} = \frac{\int_{|\vec{r}-\vec{r}_{BL}|<\sigma}\phi_s(\vec{r},t)d\vec{r}}{\int_{|\vec{r}-\vec{r}_{BL}|<\sigma}d\vec{r}}$
- Concentration of a species $s$ outside the BL = $\phi_s^{out} = \frac{\int_{|\vec{r}-\vec{r}_{out}|>\sigma}\phi_s(\vec{r},t)d\vec{r}}{\int_{|\vec{r}-\vec{r}_{BL}|>\sigma}d\vec{r}}$
- Average concentration of a species $s$ in the system = $\phi_s^{avg} = \frac{\int \phi_s(\vec{r},t)d\vec{r}}{\int d\vec{r}}$
- Protein partitioning to the BL = $\frac{\phi_s^{BL}}{\phi_s^{out}}$
- Chemical potential of species $s$ = $\frac{\delta F}{\delta \phi_s}$

#### S3 Condensate dynamics and mRNA transcription in the absence of actively transcribing lncRNAs

To get some baseline expectations, we first study the effect of actively transcribing mRNA on the condensate dynamics, in the absence of any active transcription of lncRNAs. We vary the transcription rate constant $k_M$ of mRNAs and map out the nature of the non-equilibrium steady state and the dynamics of approach. This is described in figure S5A.

As we increase the transcription rate constant $k_M$, the mRNA concentration at the BL locus at steady state increases (Figure S5B). The amount of protein at the BL locus however initially increases and then decreases (Figure S5B). This is consistent with our expectations from the re-entrant phase diagram described in section S2.3. At low $k_M$, the active transcription of mRNA which depends on the local protein concentration $\phi_P(\vec{r})$ couples with the mRNA-protein interactions to result in a positive feedback loop that helps recruit more protein to the BL. At high $k_M$, a lot of mRNA is produced in the system at steady state and there is a region in space for which it is unfavorable to form a 2-phase system. This corresponds to the case where enough mRNA is produced to locally dissolve the dense phase of protein due to re-entrant transition. From figures S5B and S5C, we can see that the protein recruitment to the BL locus goes down for high $k_M \geq 0.1$ and the dense phase of protein dissolves. It dissolves from the inside-out and the protein in the system accumulates at the periphery of the BL with most of the RNA being present in the center.

The dynamics of protein recruitment to the BL locus again has two regimes (Figure S5C, Figure S5D). At low $k_M$, increasing $k_M$ increases the recruitment of protein to the BL at steady state and results in the formation of a stable dense phase of protein (Figure S5D). At high $k_M$, the mRNA concentrations at the BL can cross $\phi_R^{BL} = \phi_R^c = 0.1$, which locally dissolves the dense phase of protein and results in a short-lived condensate (Figure S5D). This is consistent with prior experimental results relating amount of mRNA transcribed and condensate lifetimes [25].

#### S4 Supplemental figures

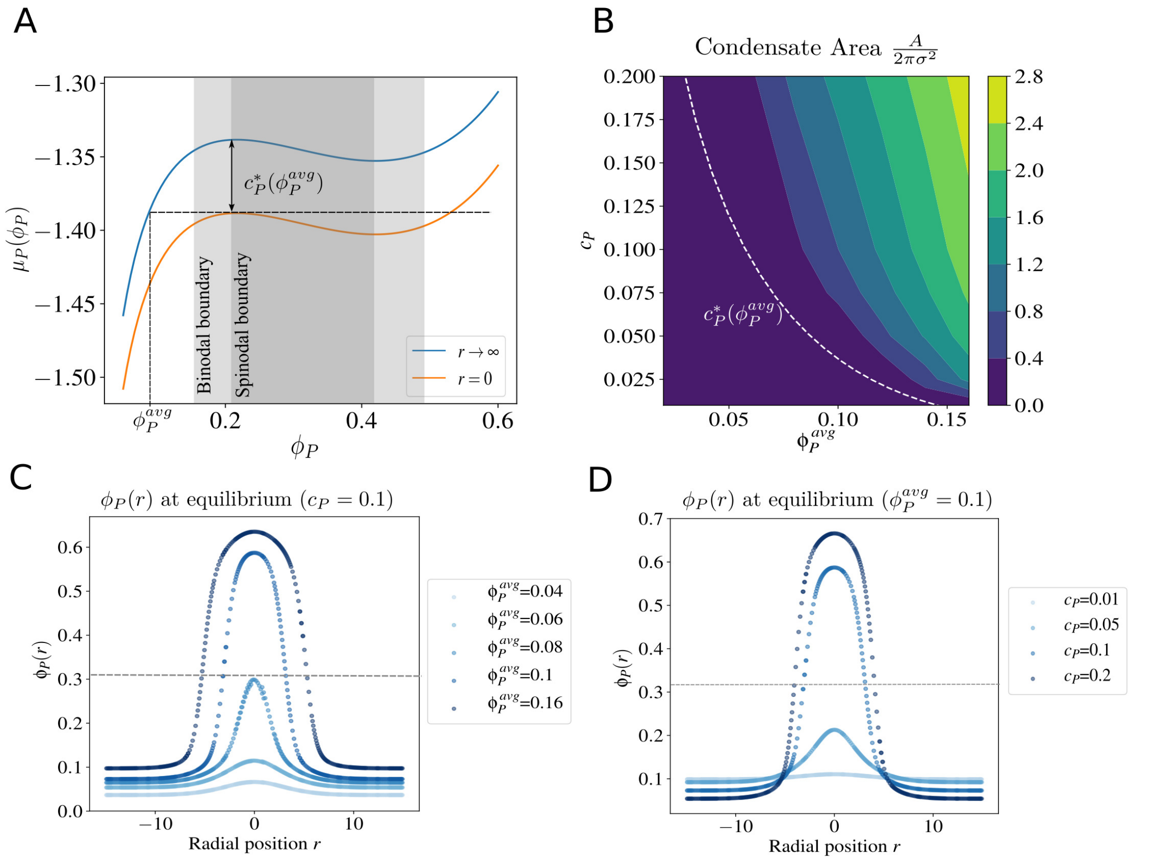

Figure S1: (A) the protein chemical potential $\mu_P$ as a function of the protein concentration $\phi_P$ for $\chi_P = 1.1$, $N_P = 5$ at $r = 0$ and $r \to \infty$. The edges of the light gray region represent the values of $\phi_P$ that correspond to the coexistence concentrations of proteins in the light and dense phase. The dark gray region represents the region of spinodal instability. For a given amount of protein in the system as quantified by $\phi_P^{avg}$, $c_P^*(\phi_P^{avg})$ is the depth of the Gaussian chemical potential well required to locally recruit enough protein at $r = 0$ to cross the spinodal boundary and form a dense phase of protein (B) Area of the transcriptional condensate $A/\pi\sigma^2$ for different amounts of protein in the system ($\phi_P^{avg}$) and depths of the chemical potential well ($c_P$). Condensate area $A$ is defined as the area of the region in space where the protein concentration $\phi_P(r) > 0.3$ (C) Profile of $\phi_P(r)$ along the radial direction at constant $c_P = 0.1$ for different amounts of protein in the system $\phi_P^{avg}$ (D) Profile of $\phi_P(r)$ along the radial direction at constant $\phi_P^{avg} = 0.1$ for different depths of the chemical potential well $c_P$

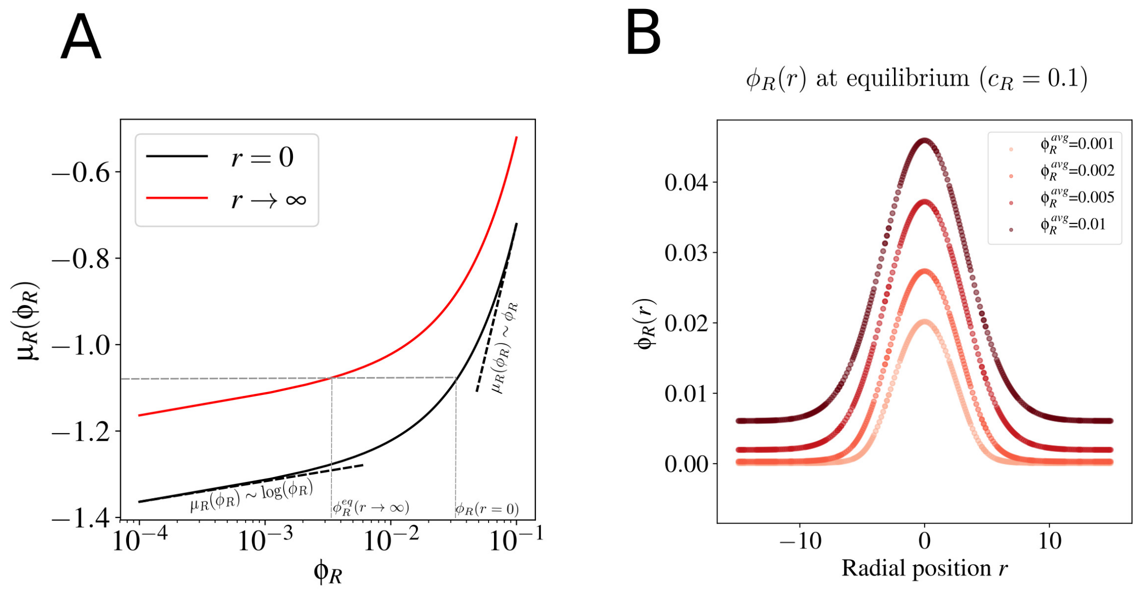

Figure S2: (A) The RNA chemical potential $\mu_R$ as a function of the RNA concentration $\phi_R$ for $c_R = 0.2$, $N_R = 50$, and $\chi_R = 2.0$ at $r = 0$ and $r \to \infty$. The RNA profile at equilibrium $\phi_R^{eq}(r)$ lies within the range $\phi_R^{eq}(r = 0)$ and $\phi_R^{eq}(r \to \infty)$, peaking at $r = 0$ (B) The equilibrium profiles $\phi_R^{eq}(r)$ for different amounts of RNA in the system as quantified by $\phi_R^{avg}$

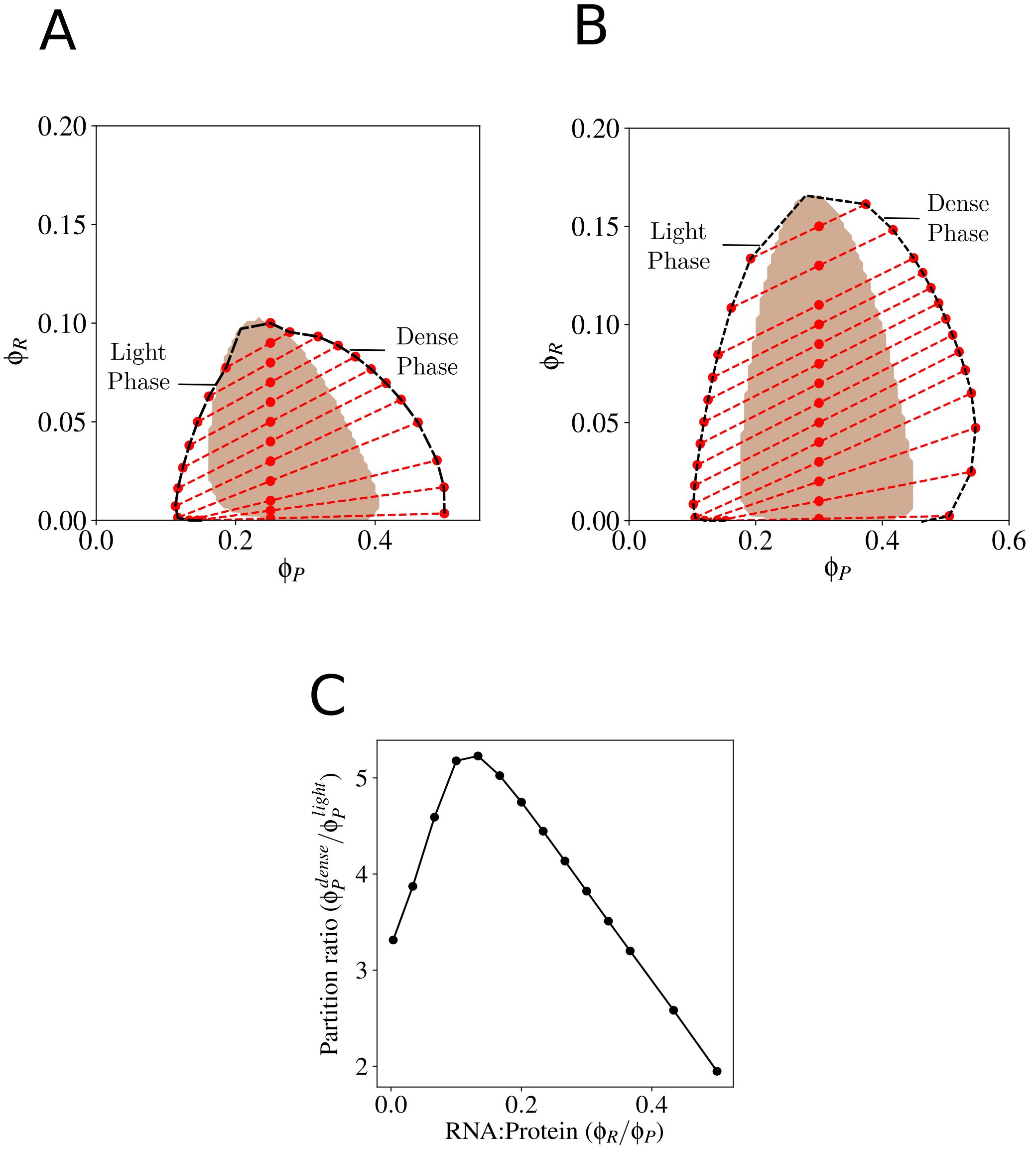

Figure S3: (A) Phase diagram for the Protein-RNA-Solvent ternary system described by $F_{FH}$ at different protein ($\phi_P$) and RNA concentrations ($\phi_R$). The parameters used were $\chi_P = 1.1$, $\chi_{PR} = 1.2$ and $\chi_R = 2.0$. The brown region is corresponds to the region of spinodal instability. The dotted red lines are coexistence curves that connect a point in the spinodal region to the dense and the light phase compositions that the system phase separates into (B) Phase diagram for the Protein-RNA-Solvent ternary system described by $F_{FH,mod}$ at different protein and RNA concentrations (C) Re-entrant phase diagram representing protein partition ratio $\phi_P^{dense}/\phi_P^{light}$ for $\phi_P = 0.3$ upon titrating the system with RNA for $F_{FH,mod}$

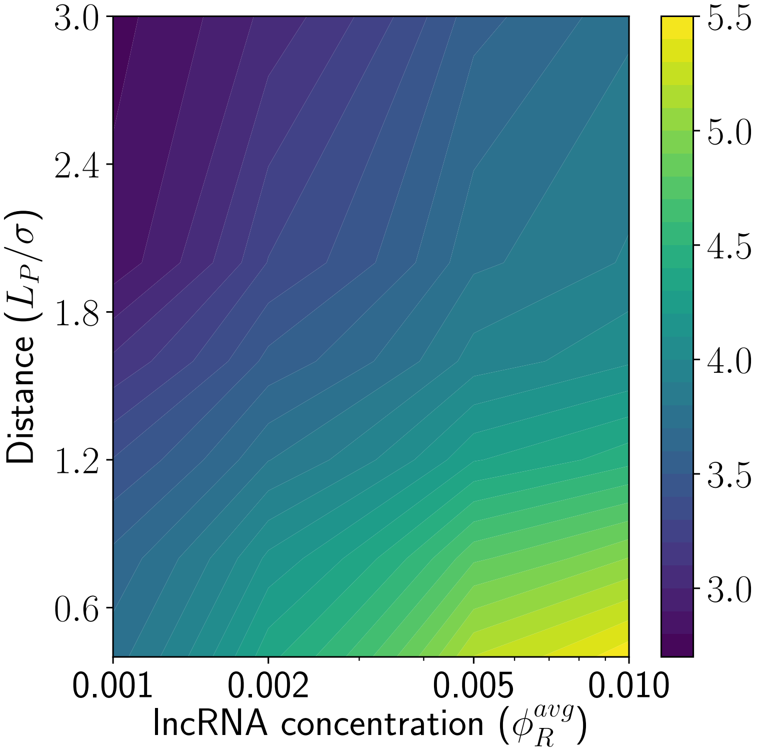

Figure S4: Phase diagram capturing how protein partitioning to the BL varies upon simultaneously varying the distance and the amount of lncRNA in the system. The light white lines indicate contours, where the effects of the lncRNA amounts and the distance can compensate each other to result in a similar protein partitioning to the BL

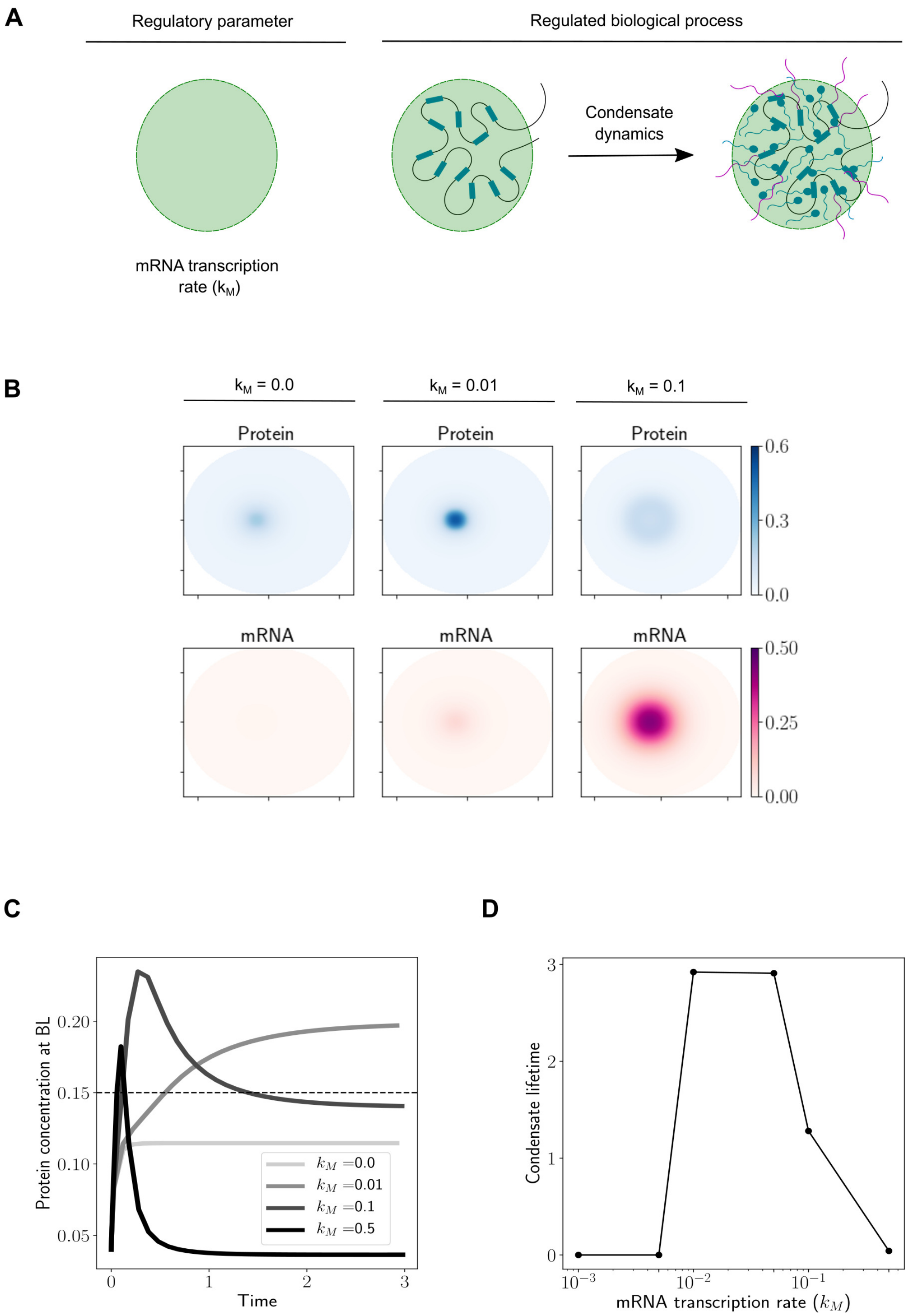

Figure S5: (A) In this figure, we will increase the mRNA transcription rate constant $k_M$ and study how that impacts protein concentration at the BL and condensate lifetimes (B) Steady state concentration profiles of protein (blue) and RNA (red) at steady state for different values of the transcription rate constant $k_M$ (E) Dynamics of protein concentration ($\phi_P^{BL}$) for different values of $k_M$. Time is in the dimensionless units of $k_d t$ (D) The dependence of condensate lifetime on $k_M$. The condensate lifetime is also reported in the dimensionless units $k_d t$, and is defined as the duration of time for which protein concentration at the BL is "appreciable". We chose a cutoff $\phi_P^{BL} > 0.15$ to define "appreciable" protein concentration at the BL. Note that this specific numerical choice of the cutoff value doesn't change the qualitative nature of the trends or results.

#### References

[1] Noa Gil and Igor Ulitsky. Regulation of gene expression by cis-acting long non-coding RNAs, 2020.

[24] Halima H Schede, Pradeep Natarajan, Arup K Chakraborty, and Krishna Shrinivas. Organization and regulation of nuclear condensates by gene activity. *bioRxiv*, 2022.

[25] Won Ki Cho, Namrata Jayanth, Brian P. English, Takuma Inoue, J. Owen Andrews, William Conway, Jonathan B. Grimm, Jan Hendrik Spille, Luke D. Lavis, Timothée Lionnet, and Ibrahim I. Cisse. RNA Polymerase II cluster dynamics predict mRNA output in living cells. *eLife*, 5(MAY2016):1–31, 2016.
